# Early childhood development of white matter fiber density and morphology

**DOI:** 10.1101/624171

**Authors:** Dennis Dimond, Christiane S. Rohr, Robert E. Smith, Thijs Dhollander, Ivy Cho, Catherine Lebel, Deborah Dewey, Alan Connelly, Signe Bray

## Abstract

Early childhood is an important period for cognitive and brain development, though white matter changes specific to this period remain understudied. Here we utilize a novel analytic approach to quantify and track developmental changes in white matter micro- and macro-structure, calculated from individually oriented fiber-bundle populations, termed “fixels”. Fixel-based analysis and mixed-effects models were used to assess tract-wise changes in fiber density and bundle morphology in 73 girls scanned at baseline (ages 4.09-7.02, mean=5.47, SD=0.81), 6-month (N=7), and one-year follow-up (N=42). For comparison, we also assessed changes in commonly utilized diffusion tensor metrics: fractional anisotropy (FA), and mean, radial and axial diffusivity (MD, RD, AD). Maturational increases in fixel-metrics were seen in most major white matter tracts, with the most rapid increases in the corticospinal tract and slowest or non-significant increases in the genu of the corpus callosum and uncinate fasciculi. As expected, we observed developmental increases in FA and decreases in MD, RD and AD, though percentage changes were smaller relative to fixel-metrics. The majority of tracts showed more substantial morphological than microstructural changes. These findings highlight early childhood as a period of dynamic white matter maturation, characterized by large increases in macroscopic fiber bundle size, mild changes in axonal density, and parallel, albeit less substantial, changes in diffusion tensor metrics.

## 1 INTRODUCTION

Early childhood is a period of rapid cognitive and behavioral maturation, with profound changes in attention, working memory, reading, math, and self-regulatory abilities (Breckenridge et al., 2013; Burnett Heyes et al., 2012; Ferretti et al., 2008; Garon-Carrier et al., 2018; Montroy et al., 2016). Maturation of these essential cognitive-behavioral skills is believed to be related to white matter growth in the brain (Bathelt et al., 2018; Jolles et al., 2016; Klarborg et al., 2013; Qiu et al., 2008; Van Eimeren et al., 2008), though developmental trajectories of white matter maturation and the biological underpinnings of these changes remain poorly understood. The reason for this is two-fold. Firstly, while white matter development throughout the lifespan has been well characterized (Lebel et al., 2012; Westlye et al., 2010; Yeatman et al., 2014), few studies have focused on white matter changes during early childhood. Secondly, most studies make use of the diffusion tensor model (subsequently referred to simply as diffusion tensor imaging or “DTI”), which provides only limited insight into the biological changes underlying white matter growth. This limitation is due to the unreliability of DTI metrics in regions of crossing/kissing fibers (Jeurissen et al., 2013) and the fact that such metrics are potentially sensitive to a range of microstructural features. Age-related changes in fractional anisotropy (FA) and mean diffusivity (MD) from birth to adulthood have been suggested to potentially reflect developmental increases in axonal packing, myelination, or so-called white matter “integrity” (Dubois et al., 2014; Lebel and Deoni, 2018; Mukherjee and McKinstry, 2006), but it is not possible to infer information related to specific tracts or to be clear which biological features are changing using these metrics. Studies utilizing higher-order diffusion models are necessary to provide a more specific description of changes in white matter structural properties during early childhood.

Advanced diffusion imaging techniques, which provide metrics with greater structural specificity, are increasingly being applied to investigate white matter development. One such technique is fixel-based analysis (FBA; (Raffelt et al., 2017b)), which is a novel approach to quantify and analyze white matter micro- and macro-structure within “fixels” (specific fiber-bundle populations within a voxel (Raffelt et al., 2015)). FBA can be utilized in combination with constrained spherical deconvolution (CSD) techniques (Dhollander and Connelly, 2016; Jeurissen et al., 2014; Tournier et al., 2007) to delineate multiple fiber bundle populations within a voxel, making it possible to calculate and assess fixel-wise metrics sensitive to axonal density/packing (“Fiber Density (FD)”), fiber bundle cross-sectional size (“Fiber Cross-section (FC)”), and a combined measure encompassing both micro- and macrostructure (“Fiber Density and Cross-section (FDC)”) (Raffelt et al., 2017b). These metrics are arguably of greater specificity and interpretability than voxel-wise DTI metrics like FA and MD (Raffelt et al., 2017b).

Studies utilizing FBA in a neurodevelopmental context suggest white matter development from late childhood to early adulthood may be driven by increases in both axonal packing/density and fiber bundle size (Genc et al., 2017b, 2018b, 2018a, 2019). Age-related increases in axonal packing/density from late childhood to adulthood have also been suggested by studies utilizing an alternative diffusion model, “neurite orientation and dispersion density imaging (NODDI)” (Chang et al., 2015; Genc et al., 2017a; Mah et al., 2017); these changes have been shown to occur alongside increasing FA and decreasing MD during late childhood to adulthood and correlate more strongly with age than DTI metrics (Genc et al., 2017a; Mah et al., 2017). Taken together, these findings suggest that increasing axonal density/packing and fiber bundle size underlie the DTI changes observed in late-childhood to adulthood, though whether this is the case in early childhood remains unclear.

Brain development is notably heterochronous, and spatial trends in white matter development during early childhood remain relatively unclear. Lifetime changes in FA and MD have been shown to occur across major white matter tracts in a posterior-to-anterior fashion, with earliest maturation of projection and commissural fibers, followed by association fibers, and then frontal-temporal fibers, which mature into adulthood (Dubois et al., 2014; Lebel et al., 2012; Westlye et al., 2010). A recent large sample study in children ages 2-8 years provides evidence in support of this pattern, showing that changes in diffusion tensor metrics (FA and MD) across early childhood are more rapid in posterior regions as compared to frontal-temporal tracts (Reynolds et al., 2019). This spatial trend is also supported by FBA (Genc et al., 2018b) and NODDI (Mah et al., 2017) studies showing that axonal density/packing changes during late childhood are greatest in the forceps major, a posterior commissural fiber. These findings suggest that developmental changes in axonal density may follow a similar spatial pattern to changes observed in DTI metrics.

The aim of the current study was to characterize developmental changes in axonal packing/density and fiber bundle size of major white matter tracts during early childhood. To do this, we utilized 3-tissue CSD (Dhollander and Connelly, 2016) in conjunction with FBA (Raffelt et al., 2017b) to assess tract-wise changes in FBA metrics (FD, FC and FDC) in typically developing girls aged 4-8 years, scanned at baseline, 6-month and 12-month follow-up. Since developmental changes in these metrics have yet to be explored in early childhood, we focus here on relatively gross-anatomical changes by investigating tract-wise growth. Only females are included as participants were recruited as controls for an ongoing study investigating brain abnormalities in a genetic disorder affecting females. However, previous literature suggests sex differences during early childhood are small (Genc et al., 2017b; Reynolds et al., 2019), suggesting our findings should generalize despite this limitation. To compare our FBA metric findings to more commonly used DTI metrics, we also assessed changes in FA and MD, as well as radial diffusivity (RD) and axial diffusivity (AD). Considering previous literature suggestive of widespread developmental increases in axonal packing and fiber bundle size from late childhood to adulthood (Chang et al., 2015; Genc et al., 2018a, 2018b, 2017a; Mah et al., 2017), we hypothesized that FD, FC, and FDC would all increase with age throughout the whole brain (but particularly in major white matter tracts), indicative of ubiquitous axonal density and cross-sectional fiber bundle growth during early childhood. Furthermore, considering evidence for varying rates of maturation across white matter regions (Dubois et al., 2014; Lebel et al., 2012; Reynolds et al., 2019; Westlye et al., 2010), we hypothesized that FD, FC and FDC changes would be most rapid in commissural and projection fibers, and slowest in frontal-temporal tracts.

## 2 MATERIALS AND METHODS

### 2.1 Participants

Only female data was available for the study as participants were recruited as control subjects in an ongoing study investigating brain abnormalities in a genetic disorder that exclusively affects females. Participants included in the current study were all typically developing with no history of psychiatric/neurological disorder or head trauma, and no contraindications to MR scanning. Seventy-six typically developing girls were recruited from the Calgary area and underwent diffusion MRI scanning and cognitive testing at baseline (ages 4.09-7.02, mean=5.47, SD=0.81). A subset also participated in 6-month (N=7, ages 4.61-7.01, mean=5.57, SD=0.77) and 12-month (N=43, ages 5.13-7.89, mean=6.43, SD=0.74) follow-up scans. Four data sets, 3 of which belonged to participants with only one time-point, were excluded from the study due to poor data quality (described in more detail below). The final sample therefore consisted of 122 total scans collected from 73 participants at baseline (N=73), 6-month follow-up (N=7), and 12-month follow-up (N=42). The study was approved by the Conjoint Health and Research Ethics Board at the University of Calgary. Informed consent was obtained from the parents and informed assent from the participants.

### 2.2 Data acquisition

Data were collected at the Alberta Children’s Hospital and included diffusion and structural (T1-weighted) MR images, as well as handedness. To acquaint participants with the scanning environment and mitigate in-scanner head motion, participants underwent a mock-scan within one week prior to MRI scanning at each time point. During the mock-scan participants practiced lying still while sounds of the MR scanning were played to them via headphones. In between the mock-scan and actual MR image acquisition, parents were asked to have their child practice lying still at home for increasing time intervals up to 18 minutes. During MRI scanning, we also used pads inserted between the participants head and the head coil on either side to reduce head motion. MRI data were acquired on a 3T GE MR750w (Waukesha, WI) scanner using a 32-channel head coil. Diffusion-weighted images (DWIs) were acquired using a 2D spin echo EPI sequence (b=2000s/mm^2^, 45 diffusion-weighted directions, 3 non-DWI (*b*=0) images, 2.5mm^3^ isotropic voxels, 45 slices, 23×23cm FOV, TE=86.2ms, TR=10s). A second set of DWIs with the same acquisition parameters was acquired at b=1000s/mm^2^ for calculation of DTI metrics. Structural images were obtained using a T1-weighted 3D BRAVO sequence (0.8mm^3^ isotropic voxels, 24×24cm FOV, TI=600ms, flip angle=10°).

### 2.3 Data and code availability

In accordance with the terms of consent given by participants, raw data utilized in this study are not available for sharing. However, processed data (tract mean values and other pertinent data) are available upon request. Code utilized for data analysis is publicly available and provided on the software packages’ websites. URLs to software packages utilized in the study are provided where applicable. This policy is in accordance with the funding bodies that supported this research as well as the Conjoint Health and Research Ethics Board at the University of Calgary.

### 2.4 Structural image processing

Structural T1-weighted images were utilized to calculate intra-cranial volume (ICV) for use as a nuisance covariate in statistical analysis. It is generally recommended that ICV be included as a covariate in FBA to account for relative differences in head size, which may vary with age and thus potentially influence morphological fixel-based white matter metrics (fiber cross-section (FC) in particular). T1-weighted images were bias field corrected (Tustison et al., 2010), and ICV estimated using the Computational Anatomy Toolbox (CAT; http://www.neuro.uni-jena.de/cat/) in SPM 12 (https://www.fil.ion.ucl.ac.uk/spm/software/spm12/).

### 2.5 DWI preprocessing and motion assessment

To mitigate effects of head motion, which are particularly prevalent in this age range, we utilized current state-of-the-art DWI pre-processing (Andersson and Sotiropoulos, 2016) provided in *FSL* (Jenkinson et al., 2012) (https://fsl.fmrib.ox.ac.uk/fsl/fslwiki), including replacement of slice-wise signal dropout (Andersson et al., 2016) and within-volume motion correction (Andersson et al., 2017). DWI volumes for which more than 20% of slices were classified as containing signal dropout were removed from the data, and scans for which more than 10% of volumes were removed were excluded from the study entirely (resulting in the exclusion of 4 scans from 3 participants). To quantify and control for the potential influence of head motion on outcome measures, we calculated the total number of signal dropout slices per dataset for use as a covariate in statistical analyses (motion characteristics of the study sample are provided in Supplementary Table 1). Following motion correction we performed default FBA preprocessing steps including; DWI bias field correction using bias fields estimated from the mean *b*=0 images (Raffelt et al., 2012b; Tustison et al., 2010), and upsampling to 1.25mm^3^, which improves contrast in DTI/CSD output images and thereby benefits downstream processing steps (Dyrby et al., 2014). These preprocessing steps were performed separately for the b=2000s/mm^2^ and b=1000s/mm^2^ diffusion shells.

### 2.6 FBA and fixel metrics

To calculate fixel-based metrics we utilized the b=2000s/mm^2^ DWIs and generally followed the *MRtrix3* (Tournier et al., 2019) (https://www.mrtrix.org/) FBA processing pipeline^1^ (Raffelt et al., 2017b), with the exception that instead of using the standard constrained spherical deconvolution approach (Tournier et al., 2007) estimation of white matter fiber orientation distributions (FODs) was done for each participant via “single-shell 3-tissue constrained spherical deconvolution” (SS3T-CSD) (Dhollander and Connelly, 2016) using a group averaged response function for each tissue type (white matter (WM), grey matter (GM) and cerebrospinal fluid (CSF)) (Dhollander et al., 2016; Raffelt et al., 2012b). This step was performed using *MRtrix3Tissue* (https://3tissue.github.io/), a fork of *MRtrix3* (Tournier et al., 2019). In brief, SS3T-CSD estimates anisotropic WM FODs from diffusion MRI data that are less influenced by isotropic GM-like and CSF-like signal contributions by separating the latter into their own model compartments. Similar to multi-shell multi-tissue CSD (MSMT-CSD; Jeurissen et al., 2014), it assumes a fixed anisotropic single-fibre WM response function and fixed isotropic GM and CSF response functions across the brain. Whereas MSMT-CSD can only estimate 3 tissue compartments for multi-shell data (Jeurissen et al., 2014), SS3T-CSD allows this for single-shell + b=0s/mm^2^ data due to a bespoke iterative estimation strategy (Dhollander and Connelly, 2016). This allows it to not have to rely on the same assumption for lower b-values (between b=0 s/mm^2^ and the largest b-value available in the data). Finally, the resulting WM FOD will also not be weighted by lower b-value(s), so the resulting apparent fibre density (FD) is less influenced by extra-axonal signal and more specifically sensitive to intra-axonal volume.

The remaining processing steps included: global intensity normalization of FOD images (Raffelt et al., 2017a); generation of a study-specific FOD template (Raffelt et al., 2011); registration of FOD images to template space (Raffelt et al., 2012a); segmentation of both template and per-scan FODs in template space to identify discrete fixels (Smith et al., 2013); and identification and assignment of corresponding fixels between the scans and the template (Raffelt et al., 2015).

Following these processing steps, we calculated the three standard fixel-based metrics for FBA (Raffelt et al., 2017b):

1. *Fiber density (FD)*: a microstructural metric that serves as a proxy for axonal density or packing;
2. *Fiber cross-section (FC)*: a macrostructural metric that approximates relative fiber bundle diameter or size;
3. *Fiber density and cross-section (FDC)*: the product of FD and FC, which encapsulates changes to both micro- and macro-structure.

These metrics are calculated from the FOD amplitude and normalization parameters in registering scan FODs to the study template space. While these metrics are proportional, they are not directly expressed in standard physical units, but in arbitrary units.

### 2.7 DTI metrics

The diffusion tensor model was applied to the preprocessed b=1000s/mm^2^ shell data to estimate FA, MD, RD and AD maps. These maps were subsequently transformed to the study-specific FOD template by first rigid-body registering the b=1000s/mm^2^ shell to the b=2000s/mm^2^ shell, and then applying the b=1000s/mm^2^-to-FOD template space transformation. b=1000s/mm^2^ shell data was not acquired for 3 participants (one first visit scan from a participant with two timepoints, and two scans from participants with only one timepoint). Data from these scans were therefore excluded from statistical analysis of DTI metrics.

### 2.8 Template-based tractography and tract means

To account for anatomical differences in the fiber bundle pathways across individuals, we extracted major white matter fiber bundles from the study-specific FOD template and utilized a single set of tracts along with normalized scan data to calculate mean tract values. A whole-brain tractogram was first generated via probabilistic tractography (Tournier et al., 2010) performed on the FOD template (Raffelt et al., 2015) using random seeding throughout the entire template, with 20 million streamlines generated (tractography parameters: step size=0.625mm, max angle between steps=22.5°, min/max fiber length=10mm/250mm, cut-off FOD amplitude=0.1). Spherical-deconvolution informed filtering of tractograms (SIFT) (Smith et al., 2013) was then used to reduce the tractogram to 2 million streamlines exhibiting reduced global reconstruction biases.

We extracted major white matter tracts of interest from this tractogram using manually-defined inclusion and exclusion ROIs (Supplementary Table 2). These tracts included: the arcuate fasciculi, cingulum bundles, corticospinal tracts, fornix bundles, inferior frontal-occipital fasciculi, inferior longitudinal fasciculi, superior longitudinal fasciculi and uncinate fasciculi, as well as the genu, body and splenium of the corpus callosum and their projections to the cortex (Figure 1).

**Figure 1.**
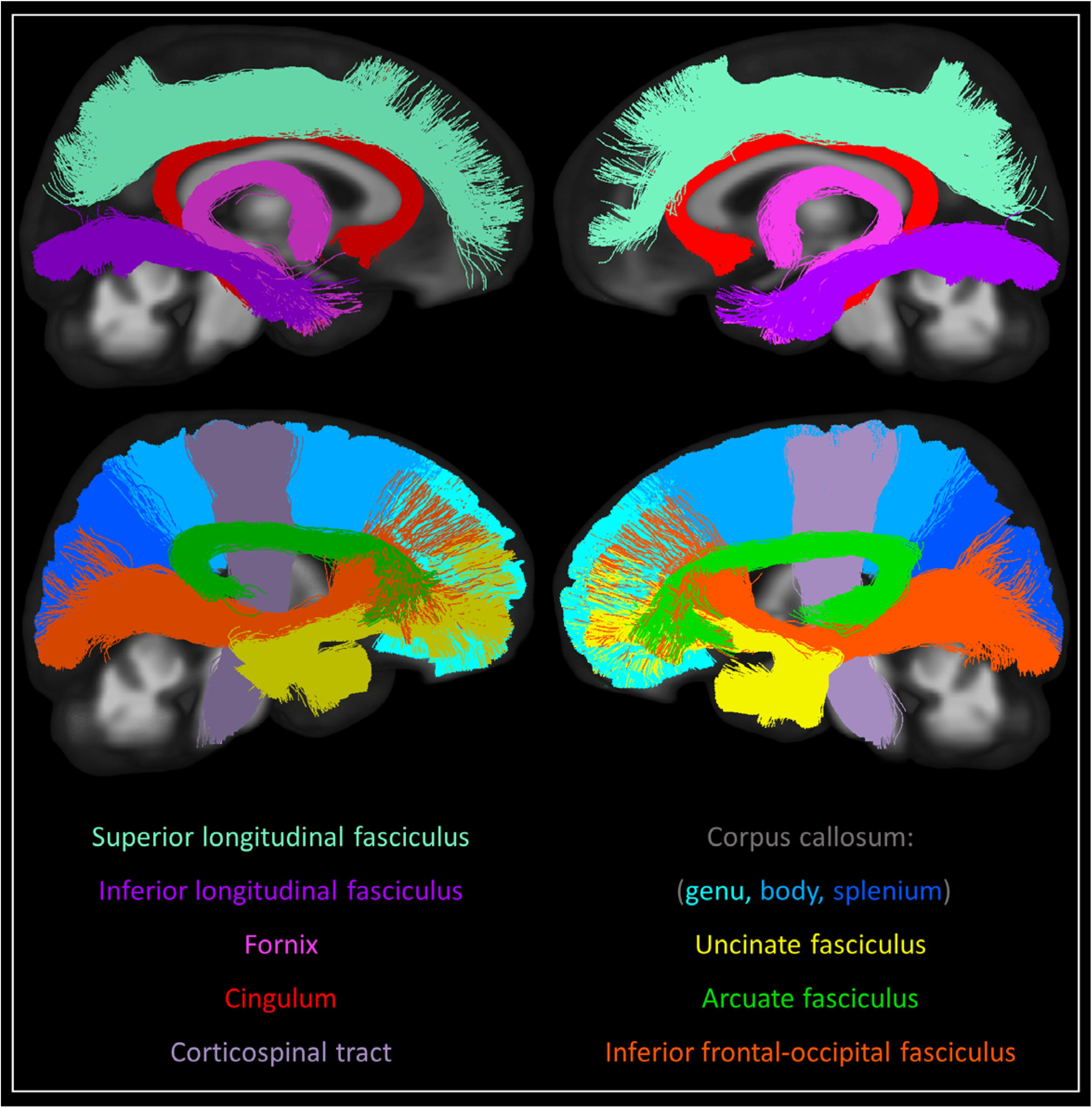
Tracts of interest. Bilateral white matter tracts of interest included in the study. Three-dimensional tracts are overlaid onto 2D slices of study template space and coloured according to the figure legend. Tracts are approximately split into medial (top row) and lateral (bottom row) groups for visual clarity.

These tracts were utilized to create both voxel and fixel masks. For each bundle, only fixels through which at least 5 streamlines passed, and that had a maximum template FOD amplitude of at least 0.1, were included in the fixel mask for that bundle; a corresponding voxel mask was then constructed that included all voxels containing at least one fixel within the respective mask.

For each scan, we calculated the mean FD, FC, FDC, FA, MD, RD and AD of each tract, by averaging values of the template-normalized metric images across voxels/fixels contained within the relevant masks (voxel masks for FA, MD, RD and AD; fixel masks for FD, FC and FDC). Additionally, to assess global brain changes, we generated global white matter voxel/fixel masks based on a maximum template FOD amplitude threshold of 0.1 and calculated mean values of voxel/fixel-wise metrics within these masks for each subject. This “global white matter mask” therefore included white matter voxels/fixels beyond those encompassed by the tract-wise masks.

### 2.9 Statistical analysis

To assess changes in tract-wise diffusion metrics with age, we utilized linear mixed effects models performed using *lme4* (Bates et al., 2017) in *R* (R Core Team, 2014). A linear mixed effects model is a regression model that can account for repeated measures on individual participants. Here, this was accomplished by modeling each participant’s intercept as a random effect. Linear mixed effects models can also account for unequal numbers of datapoints, and/or unequal time intervals between data points. This type of model is commonly utilized in a neurodevelopmental context to model maturation when there is a variable number of scans available across participants; for example, in cross-lagged longitudinal designs (Genc et al., 2019; Lebel and Beaulieu, 2011; Westerhausen et al., 2016). For all models we included age, handedness, ICV, and total number of signal dropout slices as fixed effects, and participant-specific intercepts as a random effect. Visual inspection revealed no obvious non-linear relationships or deviations from homoscedasticity or residual normality. P-values for t-statistics were obtained using the Satterthwaite’s degrees of freedom method (Hrong-Tai Fai and Cornelius, 2007), as implemented in *lmerTest* (Kuznetsova et al., 2017). False discovery rate (FDR) was utilized to control for multiple comparisons within metrics, with statistical inference set at *p*<0.05. Models were conducted with and without outliers in tract metric values, with outliers defined using quartiles (Q) and the interquartile range (IQR); the upper limit threshold for outliers was set at Q_3_ + (1.5*IQR), while the lower limit threshold was set at Q_1_ - (1.5*IQR). Exclusion of outliers did not affect inferences drawn for the results and so outlier data points were not removed in analyses reported here. We report the estimated slopes of the age regression (beta value) as the rates of change, with standard errors of the estimates as confidence intervals. For comparison of relative growth within tracts, we calculated the percent increase in a given metric by dividing the difference between the predicted value at age 8 and age 4 by the predicted value at age 4. Rates of change, predicted values at ages 4 and 8, and percent increases for each imaging metric are provided in Supplementary Tables 3-9.

Finally, to assess the relationship between metrics and changes in metric values over time, we ran Pearson correlations between FD, FC, FDC, FA, MD, RD and AD of the global white matter mask. Cross-sectional values were compared in a sample generated by selecting the scan with the least motion for each participant. To assess the relationship between longitudinal changes across metrics, we conducted Pearson correlations of the difference in metric values between baseline and 12-month follow-up scans, for those participants for whom data from both timepoints were available. Within both cross-sectional and longitudinal correlations, FDR was utilized to control for multiple comparisons, with statistical inference set at *p*<0.05. We also conducted partial correlations controlling for covariates utilized in the mixed effects models (age, handedness, ICV, and total number of signal dropout slices in cross-sectional analyses, and the difference in age between scans, handedness, change in ICV and average total number of dropout slices across scans in longitudinal correlations). Controlling for covariates, however, had a negligible affect on the results (unless otherwise stated), and we therefore report only the results from Pearson correlations without covariates.

### 2.10 Data visualization

For each imaging metric (FD, FC, FDC, FA, MD, RD and AD) we provide a figure with three panels: panel A shows tract-wise rates of change with confidence intervals to facilitate comparison of maturational rates across tracts; panel B shows linear developmental trajectories, to illustrate tract-wise developmental patterns and facilitate comparison of metric values; panel C shows an anatomical heat-map image of tract-wise rates of change, to illustrate spatial trends. Scatterplots of individual scan tract values are provided for each metric in the supplementary material (Supplementary Figures 1-5, 7, and 9). Following within-metric findings, we summarize trends in development of each tract across imaging metrics by converting tract-wise rates of change to z-scores. RD and AD are excluded from the across metric results and figure as trends in these metrics tightly followed those of MD, and their inclusion could bias across-metric conclusions and data visualization. For FD, FC, FDC, FA and MD individually, the rates of change across all regions of interest (19 tracts, as well as the global white matter mask) were converted to z-scores; thus, five z-scores were produced for each region of interest (one for FD, FC, FDC, FA and MD). Here, MD z-scores were negated to facilitate comparison of across metrics, given MD has a negative relationship with age, unlike the other metrics included in the across-metrics figure. The resulting z-scores are plotted alongside one another for visual comparison, and an anatomical image in which the five z-scores (one for each imaging metric of interest) are averaged within tracts is provided to illustrate spatial trends.

## 3 RESULTS

### 3.1 Fiber density (FD)

Significant increases in FD were seen bilaterally in the arcuate fasciculi, cingulum bundles, and corticospinal tracts, as well as the body and splenium of the corpus callosum, the right inferior frontal-occipital fasciculus, and the global white matter mask (Figure 2A-B; Supplementary Figure 1). The most rapid increases were seen in the bilateral corticospinal tract and splenium of the corpus callosum; slower statistically significant increases were seen in the body of the corpus callosum and global white matter mask; changes in the genu of the corpus callosum, fornix bundles, left inferior frontal-occipital fasciculus, inferior longitudinal fasciculus, superior longitudinal fasciculi and uncinate fasciculi were not significant. Across those tracts with statistically significant growth, FD values increased by 3.1-7.1% (Supplementary Table 3); the largest percentage increases were seen in the splenium of the corpus callosum (7.1%), the left arcuate fasciculus (6.8%), and the right cingulum (6.2%). Spatially, tracts extending into frontal cortex (i.e. the genu of the corpus callosum, inferior frontal-occipital fasciculi, and uncinate fasciculi) and temporal lobes (i.e. the inferior longitudinal fasciculi and uncinate fasciculi) showed slower and/or non-significant FD changes (Figure 2C).

**Figure 2.**
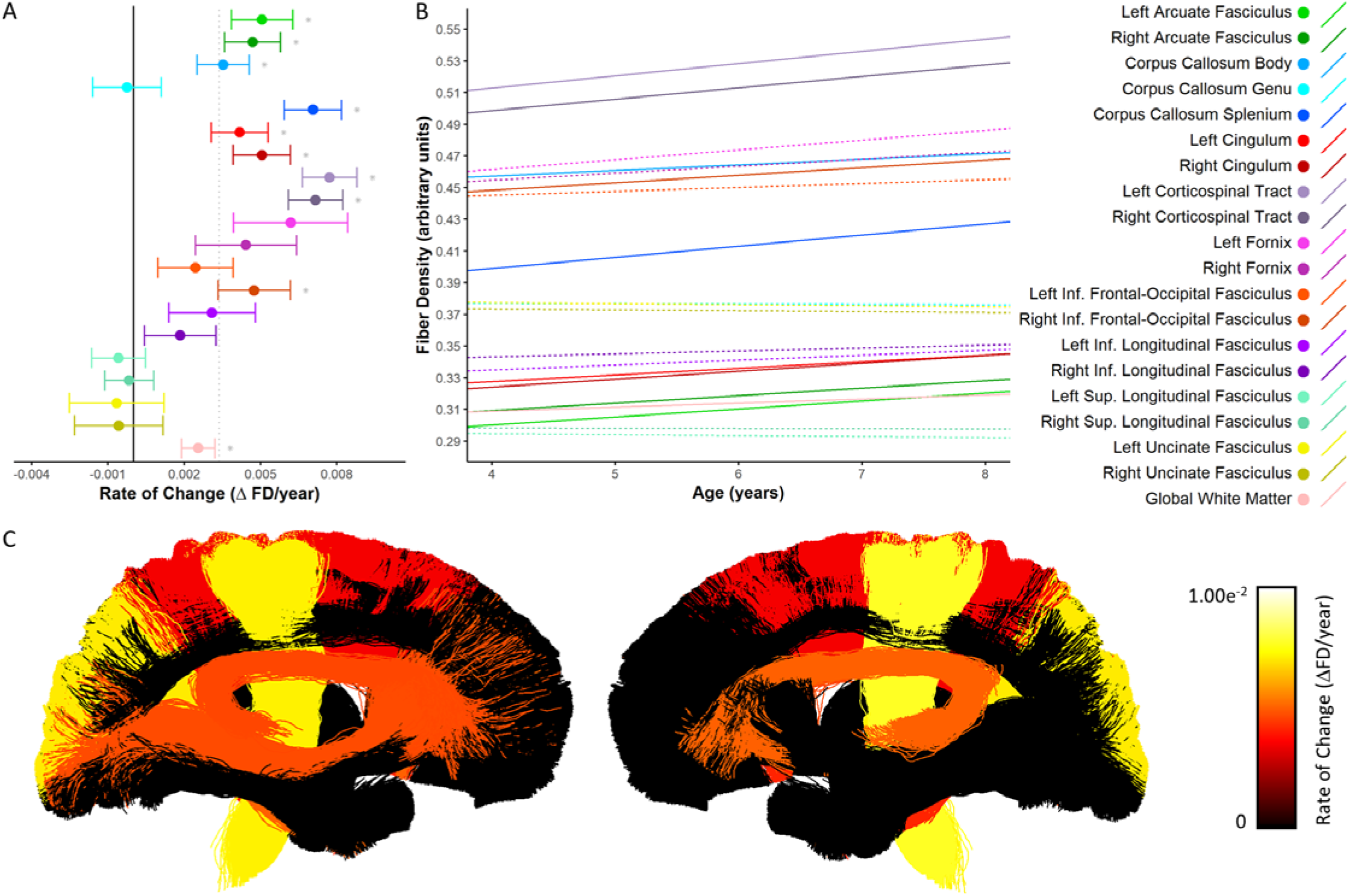
Developmental changes in fiber density. A) Rate of change in FD shown for each tract. * denotes tracts in which changes were statistically significant (p<0.05). The solid black vertical line indicates zero change, while the dotted gray vertical line indicates the mean rate of change across all tracts. B) Linear trajectories of FD growth for each tract. Solid lines indicate tracts in which there was a statistically significant relationship with age; dotted lines correspond to non-significant tracts. C) Anatomical visualization of tract-wise rates of change in FD. Streamlines corresponding to each tract are coloured according to the rate of change of FD observed in that tract. Streamlines belonging to tracts in which the rate of change was not significantly greater then zero were assigned a value of zero and appear black in the figure.

### 3.2 Fiber cross-section (FC)

Significant FC increases were seen in all white matter tracts assessed here except for the genu of the corpus callosum (Figure 3A-B; Supplementary Figure 2). Across tracts with statistically significant growth, the most rapid increases occurred in the bilateral corticospinal tract and inferior longitudinal fasciculi, while slower increases were observed in the bilateral cingulum, uncinate fasciculi, and fornix bundles. Increases ranged from 3.9-13.0% (Supplementary Table 4) across tracts with statistically significant growth. Percent increases mirrored absolute rates of change, with the greatest percentage increases in the bilateral corticospinal tract and the smallest increase in the right uncinate fasciculus. Spatially, increases were more rapid in posterior brain regions, with slower increases in tracts that connect to the frontal cortex (i.e. the uncinate fasciculi, and genu of the corpus callosum), as well as in tracts connecting deep brain structures (i.e. the cingulum and fornix bundles).

**Figure 3.**
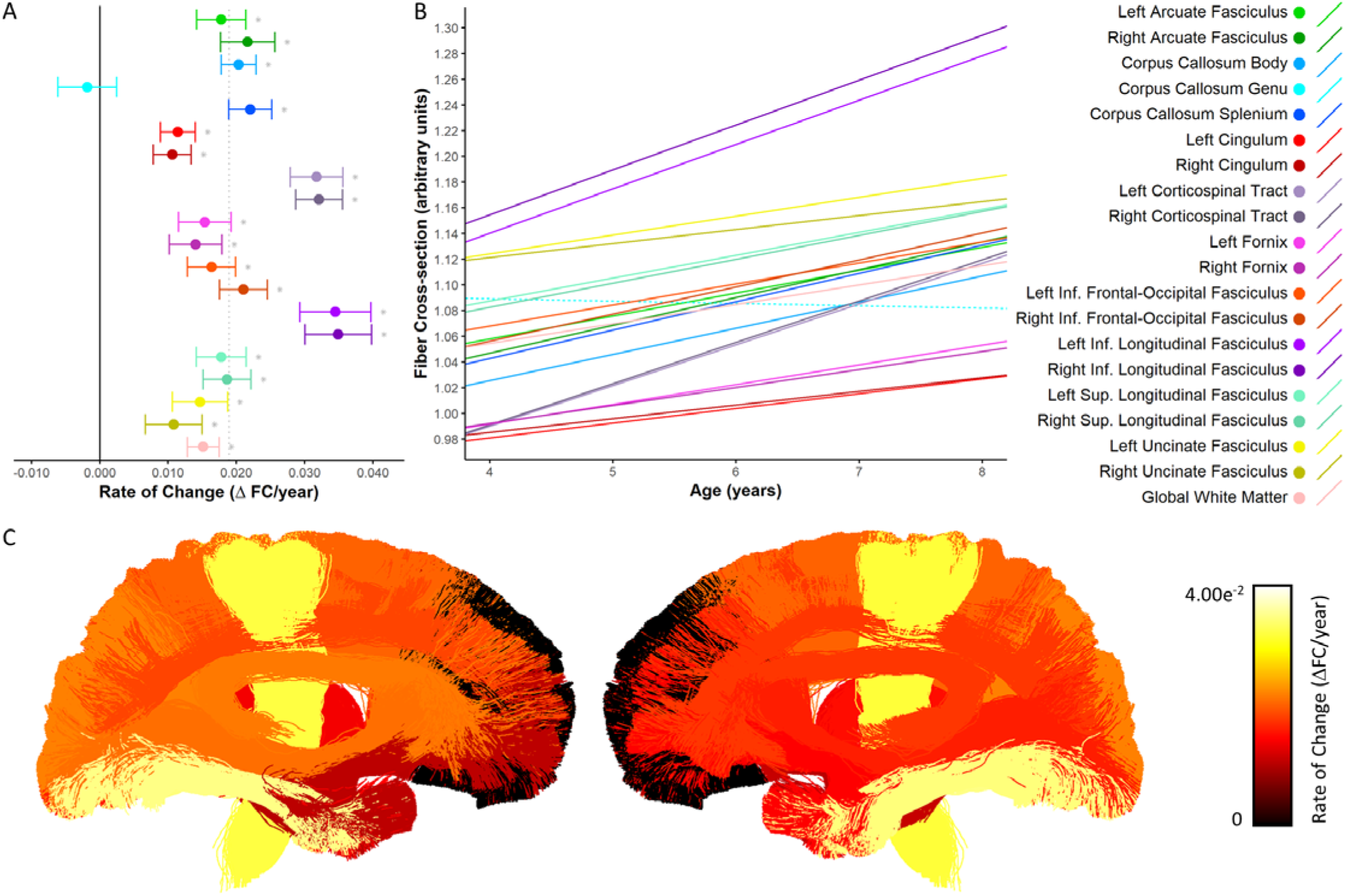
Developmental changes in fiber cross-section. A) Rate of change in FC shown for each tract. * denotes tracts in which there was significant growth. The solid black vertical line indicates zero change, while the dotted gray vertical line indicates the mean rate of change across all tracts. B) Linear trajectories of FC growth for each tract. Solid lines indicate tracts in which there was a statistically significant relationship with age; dotted lines correspond to non-significant tracts. C) Anatomical visualization of tract-wise rates of change in FC. Streamlines corresponding to each tract are coloured according to the rate of change of FC observed in that tract. Streamlines belonging to tracts in which the rate of change was not significantly greater then zero were assigned a value of zero and appear black in the figure.

### 3.3 Fiber density and cross-section (FDC)

Significant increases in FDC were seen for all tracts except for the genu of the corpus callosum, the uncinate fasciculi and the left superior longitudinal fasciculus (Figure 4A-B; Supplementary Figure 3). Across tracts with statistically significant growth, the rate of change was most rapid in the corticospinal tract bilaterally, and slowest in the right superior longitudinal fasciculus. FDC increased 6.9-21.6% across those tracts with significant growth (Supplementary Table 5), with the greatest percent increases in the corticospinal tracts, and the smallest percent increase in the right superior longitudinal fasciculus. Rate of change was generally more rapid in tracts that connect to posterior regions of the brain (i.e. the splenium and inferior longitudinal fasciculi), and slower or non-significant in tracts that make connections to the frontal cortex (i.e. the uncinate fasciculi, superior longitudinal fasciculi, and genu of the corpus callosum; Figure 4C).

**Figure 4.**
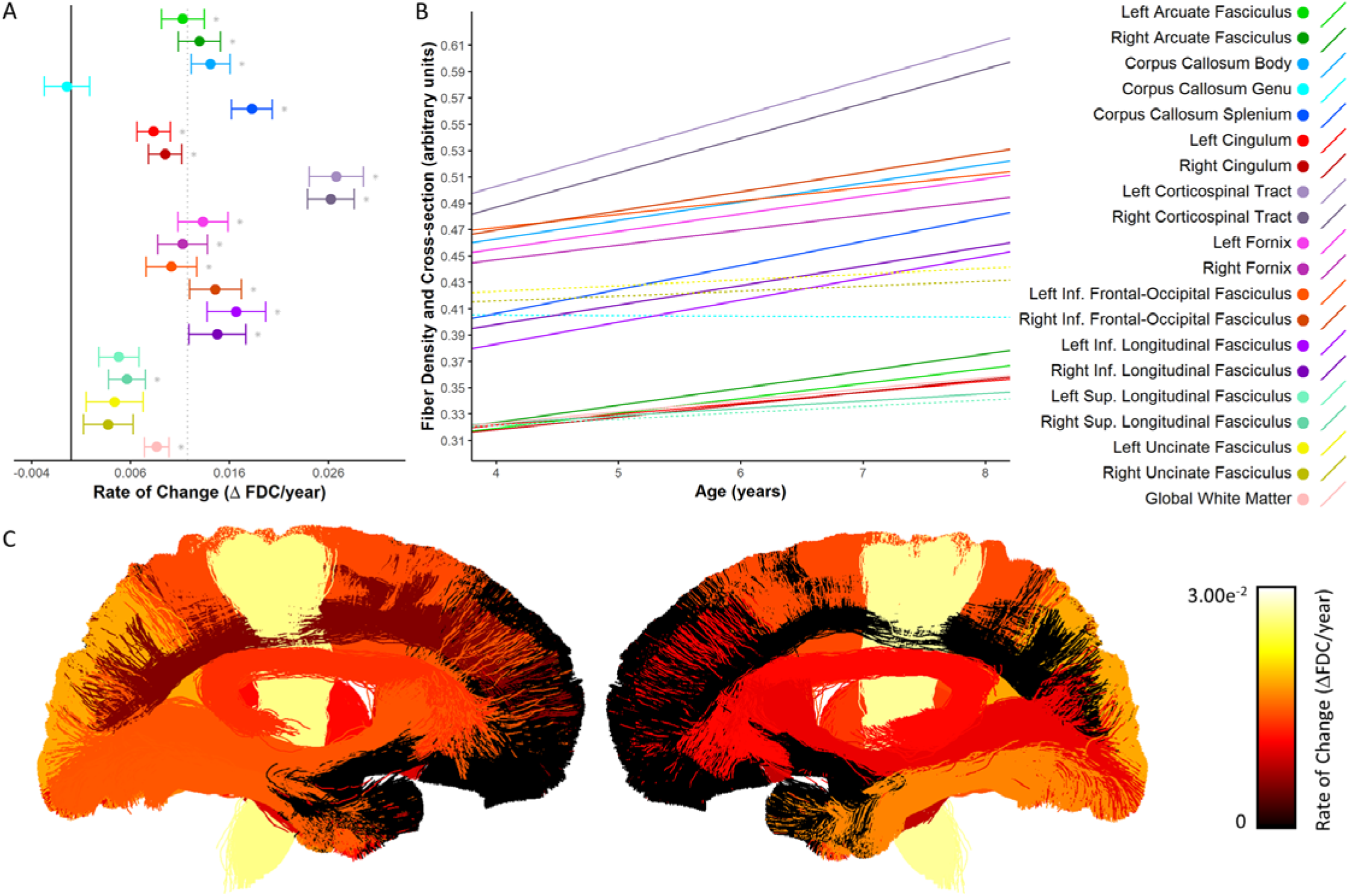
Developmental changes in fiber density and cross-section. A) Rate of change in FDC shown for each tract. * denotes tracts in which there was significant growth. The solid black vertical line indicates zero change, while the dotted gray vertical line indicates the mean rate of change across all tracts. B) Linear trajectories of FDC growth for each tract. Solid lines indicate tracts in which there was a statistically significant relationship with age; dotted lines correspond to non-significant tracts. C) Anatomical visualization of tract-wise rates of change in FDC. Streamlines corresponding to each tract are coloured according to the rate of change of FDC observed in that tract. Streamlines belonging to tracts in which the rate of change was not significantly greater then zero were assigned a value of zero and appear black in the figure.

### 3.4 Fractional anisotropy

Developmental increases in FA were observed in the arcuate fasciculi, corticospinal tracts, inferior frontal-occipital fasciculi, superior longitudinal fasciculi, splenium and body of the corpus callosum, and the global white matter mask (Figure 5A-B; Supplementary Figure 4). Amongst tracts with significant change, rates of FA increase were highly comparable across tracts; increases were slightly more rapid in the left corticospinal tract and slightly less rapid in the inferior frontal-occipital fasciculi, while changes in the genu, cingulum bundles, fornix bundles, inferior longitudinal fasciculi, and uncinate fasciculi were not significantly greater than zero. Increases ranged from 4.1-7.9% amongst tracts with significant change and were largest in the left arcuate fasciculus and smallest in the global white matter mask (Supplementary Table 6). Spatially, rates of change were faster in posterior/superior tracts, and slower in tracts that connected to anterior/inferior brain regions such as the frontal and temporal lobes (i.e. the uncinate fasciculi, genu of the corpus callosum and inferior longitudinal fasciculi; Figure 5C).

**Figure 5.**
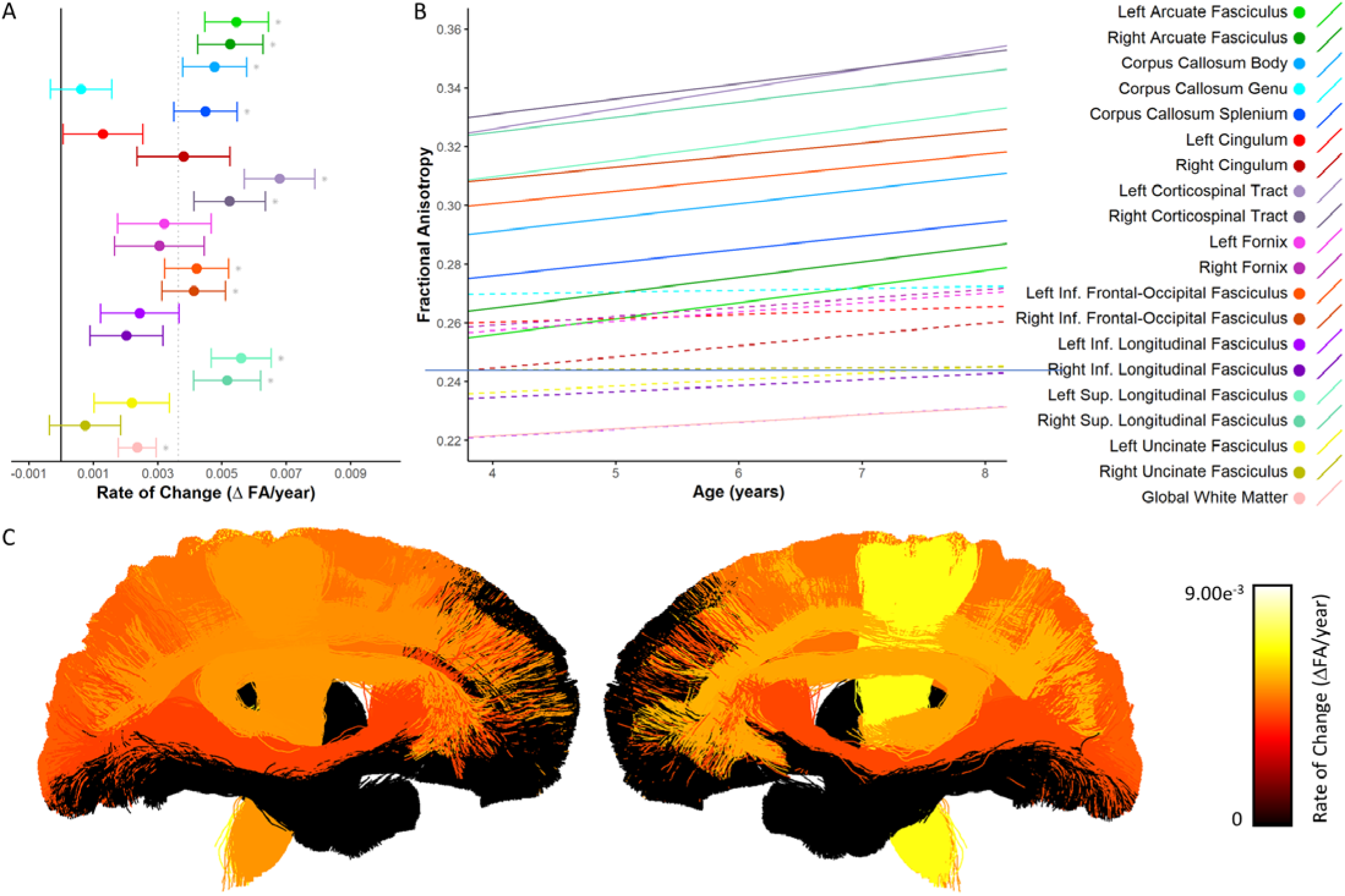
Developmental changes in fractional anisotropy. A) Rate of change in FA shown for each tract. * denotes tracts in which there was significant growth. The solid black vertical line indicates zero change, while the dotted gray vertical line indicates the mean rate of change across all tracts. B) Linear trajectories of FA growth for each tract. Solid lines indicate tracts in which there was a statistically significant relationship with age; dotted lines correspond to non-significant tracts. C) Anatomical visualization of tract-wise rates of change in FA. Streamlines corresponding to each tract are coloured according to the rate of change of FA observed in that tract.

### 3.5 Mean diffusivity

Consistent with the literature (reviewed in (Lebel and Deoni, 2018; Mukherjee and McKinstry, 2006)), significant MD decreases were observed in all examined white matter tracts (including the global white matter mask), except for in the fornix bundles (Figure 6A-B; Supplementary Figure 5). Both absolute rates of change and percent changes were highly uniform across all tracts. Decreases in MD ranged from 3.9-5.9% amongst tracts with significant change (Supplementary Table 7). Similar to other metrics reported here, MD of tracts in posterior/superior regions changed relatively faster than tracts in anterior/inferior brain regions (Figure 6C).

**Figure 6.**
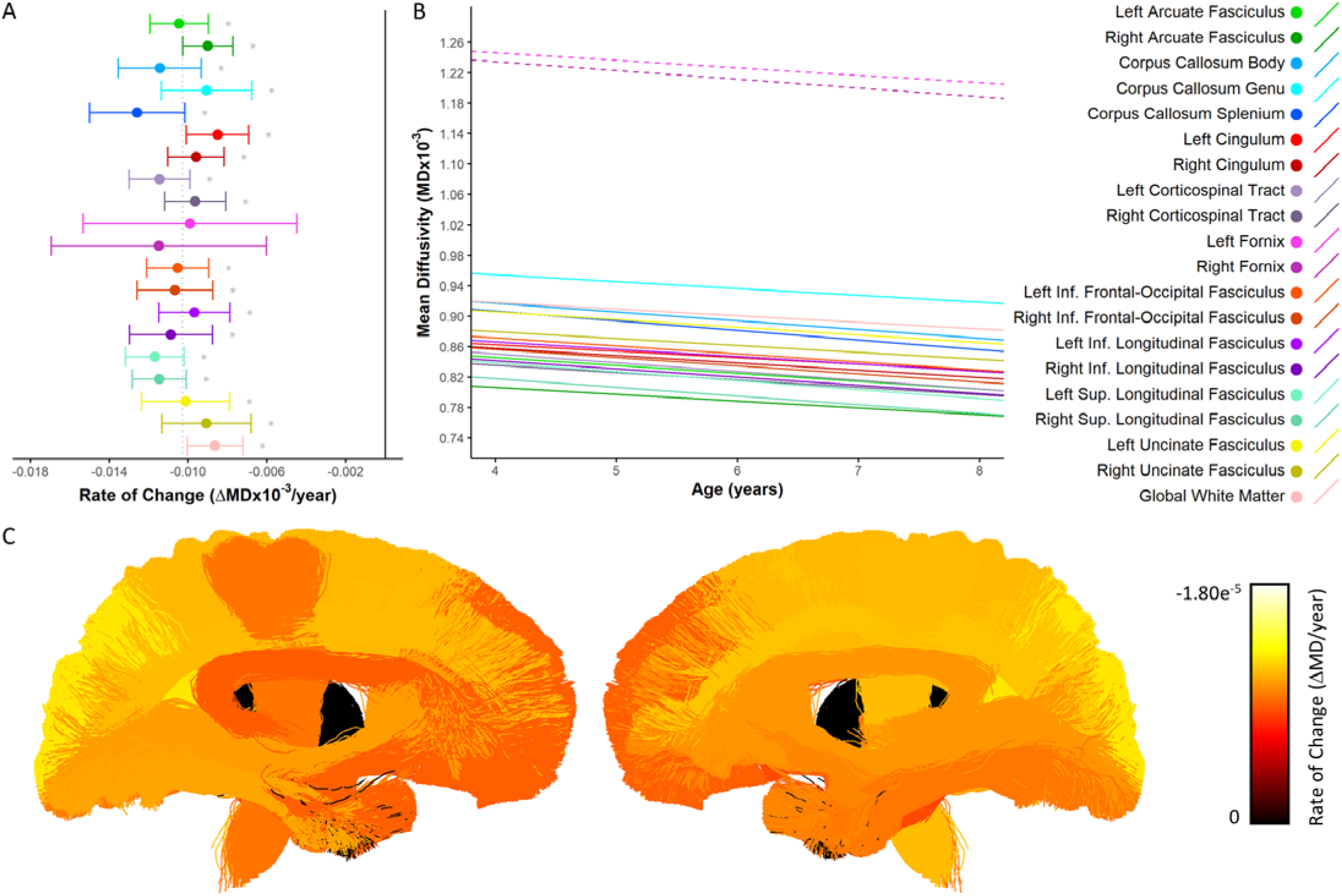
Developmental changes in mean diffusivity. A) Rate of change in MD shown for each tract. * denotes tracts in which there was significant growth. The solid black vertical line indicates zero change, while the dotted gray vertical line indicates the mean rate of change across all tracts. B) Linear trajectories of MD growth for each tract. Solid lines indicate tracts in which there was a statistically significant relationship with age; dotted lines correspond to non-significant tracts. C) Anatomical visualization of tract-wise rates of change in MD. Streamlines corresponding to each tract are coloured according to the rate of change of MD observed in that tract.

### 3.6 Radial and axial diffusivity

Changes in RD and linear trajectories essentially mirrored changes in MD, with significant decreases observed in the global white matter mask and all tracts except for the fornix bundles (Supplementary Figures 6 and 7). RD decreases ranged from 4.1-8.0% (Supplementary Table 8). Also similar to MD, rates of change were highly uniform across tracts and generally followed a posterior-anterior and superior-inferior spatial gradient of maturation. Like RD and MD, significant decreases in AD were observed in the global white matter mask and all tracts except for the fornix bundles (Supplementary Figures 8 and 9), with decreases ranging from 2.7-4.7% (Supplementary Table 9). Rates of change were highly uniform across tracts, though the genu and uncinate decreased at slightly faster rates, while AD in the arcuate fasciculi and corticospinal tracts decreased more slowly.

### 3.7 Aggregating results across metrics

Figure 7 depicts tract-wise metrics, z-scored within each metric, to facilitate visualization of maturation patterns across major FBA (FD, FC, FDC) and DTI (FA and MD) metrics. AD and RD are not included here because trends tightly followed MD. Percent increases across metrics are shown in Supplementary Figure 10. Figure 7A shows that across all metrics, more rapid changes were observed in the corticospinal tracts, and splenium of the corpus callosum, while slower change was found in the genu of the corpus callosum and the uncinate fasciculi. Tracts displayed different amounts of variability in rates of change across metrics; the inferior and superior longitudinal fasciculi had high degrees of variability, while the inferior frontal-occipital fasciculi, left corticospinal tract, and body of the corpus callosum appeared to mature more uniformly. Interestingly, for some tracts change in one imaging metric was vastly different than that of the other four (e.g. the right corticospinal tract which had particularly slow change in MD, and the inferior longitudinal fasciculi, which had particularly rapid change in FC). It is worth noting however that there was less variability in rates of change in MD compared to other metrics. In fact, in terms of absolute rates of change, the highest variability and fastest changes within a single metric was seen in FC, while the largest percent increase was seen in FDC (Supplementary Figure 10). Spatially, incorporating data across major DTI and FBA metrics, more rapid change was seen in tracts connecting to posterior/superior brain regions (e.g. the splenium of the corpus callosum and corticospinal tracts), while slower change was seen in tracts connecting to the frontal cortex (e.g. the genu of the corpus callosum and uncinate fasciculi; Figure 7B).

**Figure 7:**
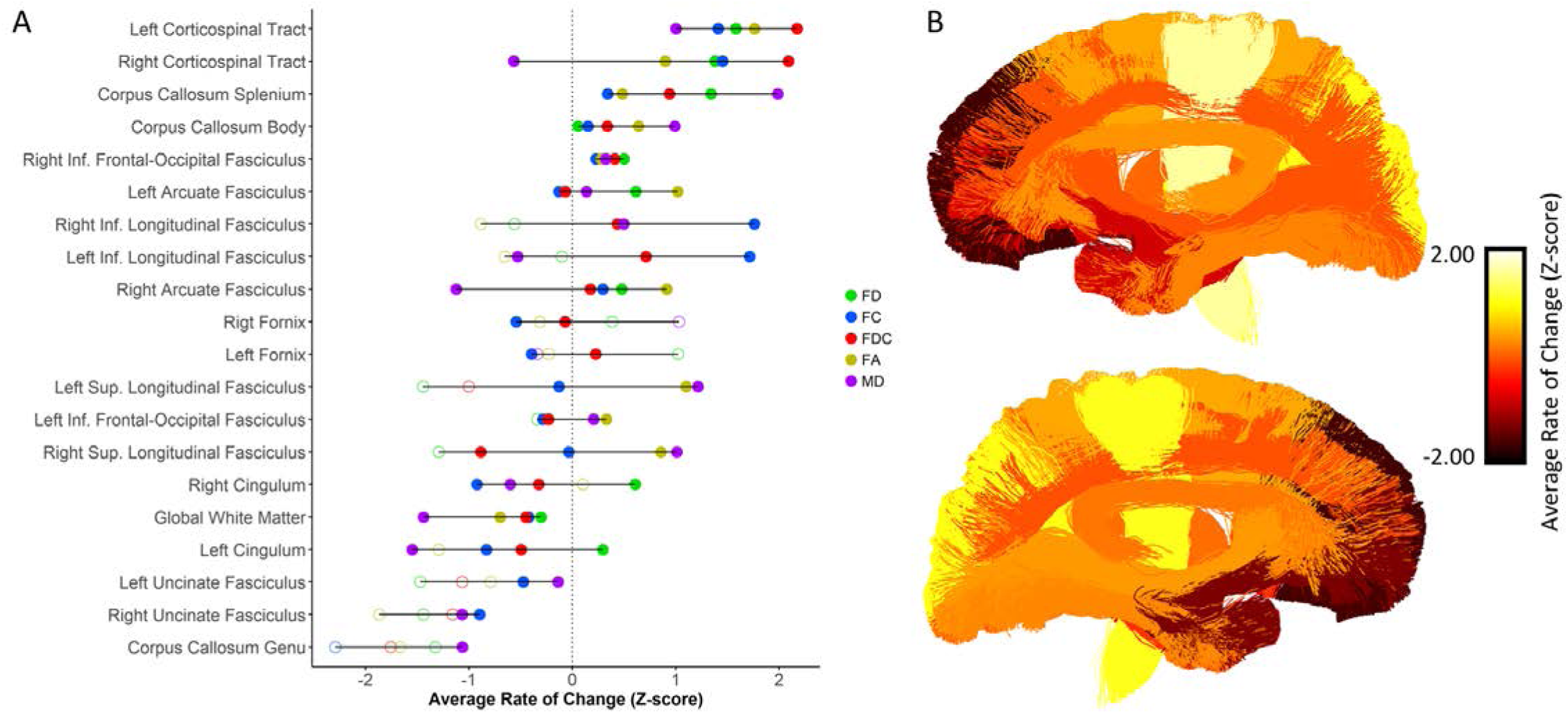
Metric-wise z-scored tract maturation. A) Tract-wise rate of change across FD, FC, FDC, FA and MD. Z-scores were calculated across tracts for each metric. Tracts are listed in descending order of average z-scored rate of change. Solid circles indicate statistically significant rates of change, while open circles indicate non-significant rates of change. The dotted gray line indicates the average rate of change across metrics (i.e. a z-score equal to zero). B) Anatomical visualization of tract-wise average z-scored rate of change across FD, FC, FDC and FA. In both panels, a negative z-score indicates a below average maturational rate, while a positive z-score indicates an above average maturational rate.

### 3.8 Correlations across metrics

Cross-sectional correlations across imaging metrics in the global white matter mask were conducted in a sample consisting of 71 participants (ages 4.22-7.39, mean=5.71, SD=0.81). There was no significant correlation between FD and FC, though unsurprisingly both correlated significantly with FDC (r=0.64 and 0.83 respectively), given FDC is the product of FD and FC. Significant correlations between FBA and DTI metrics were found for; FD and all DTI metrics (r=0.58, −0.75, −0.78, −0.61, for FA, MD, RD and AD respectively), FC and FA (r=0.52), and FDC and all DTI metrics (r=0.72, −0.29, −0.36, and −0.30 for FA, MD, RD and AD respectively). When controlling for covariates (age, handedness, total number of dropout slices, and ICV) in partial correlations, FC correlated significantly with FD (r=0.68), FDC (r=0.90), FA (r=0.60), MD (r=−0.61), RD (r=−0.64), and AD (r=−0.54). Significant correlations were also observed between all pairs of DTI metrics, with absolute r-values ranging from0.41 to 0.99. Correlation coefficients for all pairs of assessed metrics are provided in Supplementary Table 10.

A sample of 41 participants with baseline (ages 4.14-6.88, mean=5.46, SD=0.78) and 12-month follow-up (ages 5.13-7.89, mean=6.52, SD=0.75) data were utilized for correlations between changes in imaging metrics in the global white matter mask. Increases in FBA metrics were significantly correlated between all FBA metrics (ΔFD and ΔFC, r=0.59; ΔFD and ΔFDC, r=0.90; ΔFC and ΔFDC, r=0.87). Longitudinal increases in FD and FDC correlated with changes in DTI metrics; ΔFD correlated with ΔMD (r=−0.70), ΔRD (r=−0.64), and ΔAD (r=−0.63), and ΔFDC correlated with in ΔMD (r=−0.52), ΔRD (r=−0.46), and ΔAD (r=−0.51). Developmental changes in DTI metrics were significantly correlated in all cases, with absolute r-values ranging from 0.35 to 0.95. Correlation coefficients for all longitudinal correlations are provided in Supplementary Table 11.

## 4 DISCUSSION

In the current study we investigated age-related changes in fixel-based quantitative metrics and compared developmental trajectories to historically utilized DTI metrics. We found that fiber density (FD) increased in several (but not all) major white matter tracts, while fiber cross-section (FC) increased in all tracts but one. Combined micro- and macro-structural growth (fiber density and cross-section (FDC)) was observed in most tracts. These findings suggest that early childhood white matter development is characterized by relatively widespread increases in cross-sectional fiber bundle size, with additional localized increases in axonal density. These developmental changes were observed alongside significantly increasing FA and decreasing MD, RD and AD in most tracts. Importantly, tracts exhibited unique developmental trajectories within and across metrics, providing further evidence that white matter maturation during early childhood is heterochronous. Furthermore, across all metrics we observed a posterior-anterior maturational trend with faster maturation of tracts in posterior brain regions and slower maturation of frontal tracts. Below we discuss the implications of findings in FBA and DTI metrics, and compare fixel- and voxel-based results. Since findings with respect to RD and AD closely mirrored those of MD, we focus our discussion of DTI findings primarily on FA and MD.

### 4.1 FBA metrics

White matter development during childhood is believed to be primarily driven by changes in axonal packing (Lebel and Deoni, 2018). This is based in part on recent evidence that neurite density index (NDI), a proxy for axonal packing, shows widespread increases during childhood and adolescence that are associated with age more strongly than DTI metrics FA and MD (Genc et al., 2017a; Mah et al., 2017). Here we see similar regional trends to those of NDI studies, with faster FD increases in the corticospinal tract and splenium of the corpus callosum (Chang et al., 2015; Jelescu et al., 2015) and high relative changes in the cingulum bundles and arcuate fasciculi (Mah et al., 2017). Our findings are also consistent with longitudinal FBA analyses in older children (aged 9-13), showing FD increases with age in several WM tracts, including the corpus callosum, cingulum bundles, superior longitudinal fasciculi, inferior frontal-occipital fasciculi and corticospinal tracts (Genc et al., 2018b, 2019). However, our FD findings are less widespread than we hypothesized: we report no significant FD increases in the genu of the corpus callosum, fornix, inferior longitudinal fasciculi, superior longitudinal fasciculi, or uncinate fasciculi. Although we find less widespread FD changes, some are larger in magnitude than that reported in older children (3.1-7.1% as compared to 2.5-3.5% reported by (Genc et al., 2018b)). Together, these findings suggest that axonal density changes during early childhood may be characterized by profound increases in select tracts (e.g. the splenium of the corpus callosum and corticospinal tracts) and increases in frontal-temporal tracts (e.g. the uncinate fasciculi and genu of the corpus callosum) may occur at a faster rate later in childhood. It is important to note however that while FD and NDI are both believed to be sensitive to axonal packing/density, differences in our results as compared to that of previous NDI studies (Chang et al., 2015; Genc et al., 2018b, 2017a; Mah et al., 2017) may be in part due to differences in the underlying models used to derive these metrics. Further studies will therefore be necessary to validate our and these findings in relation to one another.

In addition to microstructural changes, a large number of structural (T1-weighted MRI) studies have shown developmental increases in white matter volumes from birth to adulthood (Groeschel et al., 2010; Lebel et al., 2012, 2008; Lebel and Beaulieu, 2011; Taki et al., 2013). Volumes of individual major white matter tracts have also been shown to increase with age, with each tract following a unique developmental trajectory (Lebel et al., 2012; Lebel and Beaulieu, 2011). Tract volume (Brouwer et al., 2012; Taki et al., 2013) and fiber cross-section (Genc et al., 2018b, 2019) have been shown to increase during childhood in most major white matter tracts, with slower increases in the genu of the corpus callosum and uncinate fasciculi (Genc et al., 2018b; Taki et al., 2013); findings that are consistent with our FC results. Our group has also previously shown, in an independent dataset, directional selectivity in age-associations of white matter volumes: a superior-inferior oriented white mater region encompassing the corticospinal tracts showed the largest age-slope in early childhood (Bray et al., 2015). This is consistent with the rapid FC changes in the CST reported here. Overall, our FC results suggest that early childhood is a particularly important period for development of the inferior longitudinal fasciculi and corticospinal tracts; while slower maturation in the uncinate fasciculi and superior longitudinal fasciculi, and no significant FC changes in the genu of the corpus callosum, may suggest later maturation of these tracts. It is also worth noting that FC changes were both more widespread, and showed larger percentage increases, than changes in FD. This suggests that while concurrent increases in axonal packing and fiber cross-section are prominent during early childhood (as evident from the large percentage changes observed in FDC), increases in fiber bundle size may play a relatively larger role in white matter maturation during this period. A model for white matter development in which changes in fiber density and fiber cross-section occur concurrently but via separate trajectories and mechanisms is supported by our finding of only a moderate correlation between longitudinal increases in FD and FC.

### 4.2 DTI metrics

To compare our FBA results to that of commonly utilized DTI metrics, we additionally assessed developmental changes in FA, MD, RD and AD as part of this study. Unsurprisingly, we observed developmental increases in FA and decreases in MD in most major white matter tracts; findings that are consistent with a large body of literature (see (Lebel and Deoni, 2018; Mukherjee and McKinstry, 2006) for review). Consistent with a recent DTI study in early childhood (Reynolds et al., 2019), we also found decreasing AD and RD with age in most white matter tracts. Tract-wise decreases in RD were slightly faster than that of AD, suggesting that both contributed to decreases in MD while the imbalance between the two drove changes in FA. It is important to note however, that while AD and RD have historically been viewed as metrics that provide further insight into structural properties that underlie change in white matter anisotropy, these metrics (and voxel-averaged metrics more generally) are inherently difficult to interpret in voxels with multiple fiber bundles, which make up ∼90% of white matter voxels in the brain (Jeurissen et al., 2013). These results therefore need to be interpreted with caution.

Across the lifespan, tract maturation as measured by DTI metrics has been shown to occur earliest for commissural and projection fibers, followed by association fibers, and then frontal-temporal projections (Dubois et al., 2014; Hermoye et al., 2006; Lebel et al., 2012; Lebel and Beaulieu, 2011; Westlye et al., 2010; Yoshida et al., 2013). This is broadly consistent with our results: for FA we find that changes are more rapid in projection fibers (i.e. the corticospinal tract) and slower in frontal-temporal fibers, with no significant growth in bundles such as the uncinate fasciculi and genu; for MD, RD and AD we find that commissural projections change slightly more rapidly (e.g. the splenium of the corpus callosum), and frontal-temporal fibers mature at slightly slower rates (e.g. the uncinate fasciculus and genu for MD and RD, and the arcuate fasciculi for AD).

Our results are also broadly consistent with studies that have investigated DTI metric changes from early childhood to early adulthood (Reynolds et al., 2019; Taki et al., 2013), though here we note some differences. A large DTI study in children aged 2-8 years similarly found slower FA and MD change in the uncinate fasciculi and genu of the corpus callosum (Reynolds et al., 2019), and no significant MD change in the fornix bundles. However, in contrast to our findings, they found that FA change of the corticospinal tract and splenium of the corpus callosum was slower than most other tracts, while changes in the inferior longitudinal fasciculi were most rapid. This may be attributed to the wider age range of participants in their study, or sex differences in white matter development (as our study included females only). Rapid changes in DTI metrics in the corticospinal tracts (Taki et al., 2013) and splenium (Brouwer et al., 2012) have been shown in early childhood to early adulthood, which might suggest that maturation in these tracts is more rapid in mid-late childhood than in very early childhood.

### 4.3 FBA vs DTI metrics

Unsurprisingly, we observed overlapping developmental changes in FBA and DTI metrics, suggesting that the underlying white matter maturation leads to observation of similar spatial patterns whether inferred from fixel-based quantitative measures or diffusion tensor-derived parameters. Across metrics, maturation followed a posterior-to-anterior spatial trend, with rapid maturation in the corticospinal tracts and splenium, and slower maturation in the genu of the corpus callosum and uncinate fasciculi. These findings are consistent with our hypothesis regarding spatial trends in tract-wise maturation, and are consistent with trends reported in the literature (Dubois et al., 2014; Lebel et al., 2012; Reynolds et al., 2019). Interestingly, however, tracts displayed varying degrees of uniformity in maturational rates across all metrics: the inferior longitudinal fasciculi and superior longitudinal fasciculi had large degrees of variability, while the body of the corpus callosum, right inferior frontal-occipital fasciculus and corticospinal tracts matured at a relatively consistent rate across metrics. This might suggest multifaceted structural maturation is more coupled in some tracts than others, though this has yet to be explicitly tested and the implications of this remain unclear. Future investigations into the potential mechanisms for coupling of white matter structural maturation and how concurrent and non-concurrent growth influences structural connectivity and brain function will be of high interest.

Despite spatial similarities, we did observe important distinctions in developmental trends as measured by changes in FBA and DTI metrics: percentage changes in tensor metrics were both less variable between tracts, and smaller in magnitude, than those observed in fixel-based metrics. These findings suggest that changes in non-specific, voxel-aggregate metrics inferred from the tensor model are smaller in magnitude and more homogenous across tracts during early childhood than changes in fixel-specific metrics. Interestingly, while we did observe significant correlations between fixel- and voxel-based metrics, as well as correlations between longitudinal increases, the average absolute value of correlation coefficients for fixel-voxel metric associations was relatively low (0.44 in cross-sectional brain metric correlations and 0.36 for longitudinal increases). This suggests that while FBA and DTI metrics are likely sensitive to some overlapping microstructural properties, these metrics provide complementary information, or at the very least have different sensitivity profiles. Not surprisingly, we also observed stronger correlations amongst DTI metrics than amongst FBA metrics, which is likely attributed to voxel-averaging and the inability of tensor metrics to capture fiber bundle-specific properties. Taken together these observations might suggest that FBA is better suited to tracking developmental changes in white matter maturation during early childhood, and may in turn be better able to identify white matter abnormalities in individuals with neurodevelopmental disorders (Dimond et al., 2019). It is important to note, however, that while DTI metrics have been linked to histological white matter properties (Seehaus et al., 2015), the direct link between FBA metrics and cellular properties has not been robustly validated. Future studies should examine the underlying biological properties measured by fixel-based metrics and investigate their microstructural sensitivity and anatomical specificity (pertaining to fiber bundle populations) in comparison to voxel-based DTI metrics.

### 4.4 Cognitive-behavioral and functional relevance

Early childhood is a period of profound maturation of cognitive and behavioral skills, including attention (Breckenridge et al., 2013; Mullane et al., 2016), working memory (Burnett Heyes et al., 2012), reading (Ferretti et al., 2008), math (Garon-Carrier et al., 2018), and self-regulation (Montroy et al., 2016). These skills have been linked to white matter properties during childhood (Bathelt et al., 2018; Jolles et al., 2016; Klarborg et al., 2013; Qiu et al., 2008; Van Eimeren et al., 2008), though the relationship between cognitive-behavioral skills and axonal density and fiber bundle size remains largely unexplored. One study has shown an association between FD and inattentive/internalizing behavior in typically developing children (Genc et al., 2019), while we have previously reported an association between FD and social skills in adolescents and young adults with Autism Spectrum Disorder (Dimond et al., 2019). These findings open the possibility that fixel-metrics might explain cognitive capabilities in healthy individuals as well as relative deficits in individuals with neurodevelopmental disorders. Here we found that FD, FC, and FDC increases were rapid in projection and commissural fibers (the corticospinal tracts and splenium), moderate in limbic tracts (the fornix and cingulum bundles), and slow or insignificant in frontal tracts (the genu, superior longitudinal fasciculi and uncinate fasciculi). These maturational trends may be related to rapid development of sensorimotor processes (Aboitiz and Montiel, 2003; Welniarz et al., 2017), ongoing development of emotional regulation (Ahmed et al., 2015), and protracted development of executive functions (De Luca and Leventer, 2011), respectively. We also observed rapid increases in FC of the inferior longitudinal fasciculi, which may be related to higher-order visual, emotional and reading capabilities (Herbet et al., 2018).

The mechanism by which individual differences in FD and FC might relate to cognitive-behavioral differences is unclear, though this is possibly mediated through influence on network connectivity in the brain. Increased axonal packing and fiber bundle size likely contribute to increases in structural connectivity over the course of neurodevelopment (Dennis et al., 2013; Hagmann et al., 2010), which may be related to developmental increases in functional network connectivity (Bennett and Rypma, 2013; Hagmann et al., 2010), though the relationship between these metrics likely varies across the brain (Vazquez-Rodriguez et al., 2019). Work from our research group has demonstrated that functional network connectivity is strongly associated with age during early childhood (Rohr et al. 2017, 2018; Long et al. 2017). We have also previously shown that such functional connectivity is associated with cognitive skills such as attention in the same sample of participants as used in the current study (Rohr et al., 2019, 2018, 2017). Together, this research provides a potential theoretical framework to explain the interplay between structural, functional and cognitive development. Future longitudinal studies will hopefully provide insight into the relationship between fiber density and fiber cross-section changes during early childhood, developmental increases in structural and functional brain connectivity, and cognition-behavioral maturation in typical and atypical neurodevelopment.

### 4.5 Strengths and limitations

Strengths of the current study include a relatively large sample size, and mitigation of potential confounding effects of motion through careful data screening, correction during preprocessing, and regression of motion in statistical analyses. Though we utilized a slightly lower b-value than is generally recommended for CSD (Tournier et al., 2013) and quantification of fiber density (Raffelt et al., 2012b), our study made use of state-of-the-art “single-shell 3-tissue CSD” (SS3T-CSD; (Dhollander and Connelly, 2016)), which is capable of isolating the diffusion signal attributable to white matter by modelling and removing the contribution from grey matter and CSF (Dhollander et al., 2017). Furthermore, though we acquired the minimum number of diffusion directions recommended for CSD, we were still able to achieve a sufficiently high maximum spherical harmonic degree to fully capture all content of the diffusion weighted signal for all DWIs (Tournier et al., 2013).

One limitation of the current study is our exclusive use of female participants, which blinds us to any potential sex differences in white matter development. The data utilized here were obtained from control participants in an ongoing study investigating brain abnormalities in girls with Turner syndrome, a disorder that only affects females; hence typically-developing control participants recruited for this study were correspondingly exclusively girls. While evidence for sex differences in white matter maturation is mixed (see (Lebel and Deoni, 2018) for recent review), some studies point towards small but significant sex difference in DTI (Reynolds et al., 2019) and FBA (Genc et al., 2017b) metrics. Additionally, while we tested exclusively for linear trends in white matter maturation, a recent DTI study has shown that FA and MD of select tracts follow quadratic trajectories over early childhood (Reynolds et al., 2019). Given the narrow age and limited number of scans (1-3) for each participant in our study, our sensitivity to non-linear trends would have been limited. Another potential limitation of our study is that we could not assess how variability in motion characteristics across diffusion shells may have differentially affected our FBA and DTI metric analyses. DTI conventionally uses lower b-value data than is required for an FBA. So that our DTI results would be comparable to those of other studies and consistent with conventional practice in the field we utilized b=1000s/mm^2^ DWIs for DTI estimation, though b=20000s/mm^2^ DWI were utilized in our FBA. While we controlled for motion in our analyses, differences in motion characteristics across DWI shells could potentially have influenced our comparison of FBA and DTI metric results. Lastly, while we focus here on tract-wise trends to provide insight into how fiber density and cross section change at a gross-anatomical level, maturational patterns may differ regionally along a fiber bundle. At the time of writing, mixed effects modelling at the level of individual fixels is not yet supported by *MRtrix*, though alternative fixel-level statistics could be used by future studies to investigate regional growth trends.

### 4.6 Conclusion

Here we report widespread developmental increases in tract-wise fiber cross-section between the ages of 4-8 years, with additional increases in fiber density within specific tracts. Combined micro- and macro-structural growth (FDC) demonstrated the greatest percentage changes over this age range, while fixel-based metrics in general showed both greater percentile changes, and greater between-tract variability, than voxel-based DTI metrics. These findings highlight early childhood as a period of dynamic white matter maturation, characterized primarily by developmental changes in fiber bundle size, with moderate changes in axonal packing density, and ongoing, albeit less pronounced changes in DTI properties. Our results add to the understanding of, and help to bridge gaps in, the characterization of white matter development from birth to adulthood and demonstrate the utility of FBA in this neurodevelopmental context.

## ACKNOWLEDGEMENTS

We thank all the families who participated in this study, as well as staff at the Alberta Children’s Hospital Imaging Centre.

## Footnotes

1 https://mrtrix.readthedocs.io/en/3.0_rc2/fixel_based_analysis/mt_fibre_density_cross-section.html

