## Supplementary for "Early childhood development of white matter fiber density and morphology" (from another version of this preprint: version 2, 2019-10-31)

**SUPPLEMENTARY MATERIAL**

| | | Between-Volume Motion | | | Within-Volume Motion | | |
|---|---|---|---|---|---|---|---|
| | TDS | Avg. translation (mm) | Avg. rotation (degrees) | Avg. motion (mm) | Avg. translation (mm) | Avg. rotation (degrees) | Avg. motion (mm) |
| **b=1000s/mm²** | 10.723 | 0.436 | 0.275 | 0.338 | 0.057 | 0.089 | 0.135 |
| **b=2000s/mm²** | 21.746 | 0.817 | 0.377 | 1.146 | 0.064 | 0.105 | 0.156 |

**Supplementary Table 1. Motion metrics across the study sample.** Motion metrics for the b=1000s/mm² and b=2000s/mm² diffusion weighted images of all scans included in the study. TDS=Total dropout slices. Average motion was calculated for each scan as the average translation and rotation, after converting rotation from degrees to millimetres about a 50-millimetre sphere.

| | Inclusion region of interest | Exclusion region of interest |
|---|---|---|
| **Arcuate fasciculus** | Corona radiata: medial to the superior circular/insular sulcus – coronal plane ROI<br><br>Inferior frontal cortex; medial to Broca's area – sagittal plane ROI<br><br>Temporal-parietal junction – sagittal plane ROI | Inferior temporal lobe – volumetric ROI<br><br>Prefrontal cortex – volumetric ROI |
| **Cingulum** | Anterior cingulate cortex: anterior to the genu of corpus callosum – axial plane ROI<br><br>Anterior cingulate cortex: superior to genu of the corpus callosum – coronal plane ROI<br><br>Medial cingulate cortex: superior to the body of the corpus callosum – coronal plane ROI<br><br>Inferior cingulate cortex: superior to the splenium of the corpus callosum – coronal plane ROI<br><br>Dorsal cingulate cortex: posterior to the splenium of the corpus callosum – axial plane ROI | Prefrontal cortex – volumetric ROI<br><br>Medial occipital lobe – volumetric ROI |
| **Corticospinal tract** | Cerebral peduncles at midbrain level – axial plane ROI<br><br>Precentral gyrus/primary motor cortex – axial plane ROI | Midbrain medial to the cerebral peduncles – axial plane ROI<br><br>Post central gyrus – axial plane ROI |
| **Fornix** | Columns of the fornix; superior to the anterior commissure – axial plane ROI | Midbrain and pons – volumetric ROI<br><br>Splenium of the corpus callosum and occipital lobe – volumetric ROI |

| | | |
|---|---|---|
| | Body of the fornix; superior to the thalamus, inferior to the tail of the septum pellucidum – coronal plane ROI<br><br>Posterior pillars of the fornix; posterior to the thalamus, anterior to the splenium of the corpus callosum – axial plane ROI | |
| **Inferior frontal occipital fasciculus** | Frontal cortex white matter; inferior to the claustrum and putamen – coronal plane ROI<br><br>Medial temporal white matter; inferior to the retroinsular cortex – coronal plane ROI<br><br>Inferior occipital lobe – coronal plane ROI | Inferior temporal lobe – axial plane ROI<br><br>Superior frontal and parietal lobes – volumetric ROI<br><br>Superior external capsule and corpus callosum – volumetric ROI |
| **Inferior longitudinal fasciculus** | Temporal pole – volumetric ROI<br><br>Inferior temporal lobe – coronal plane ROI<br><br>Anterior occipital lobe – coronal plane ROI | Superior frontal and parietal lobes – volumetric ROI |
| **Superior longitudinal fasciculus** | Corona radiata: superior to the head of the caudate nucleus – coronal plane ROI<br><br>Corona radiata: below precentral sulcus – coronal plane ROI<br><br>Corona radiata: below postcentral sulcus – coronal plane ROI | Temporal lobe – volumetric ROI<br><br>Inferior occipital lobe – volumetric ROI<br><br>Basal ganglia – volumetric ROI |
| **Uncinate fasciculus** | Frontal lobe white matter; superior to the short insular gyrus, inferior to the ventral claustrum – coronal plane ROI<br><br>Inferior temporal lobe – axial plane ROI | Regions posterior to the mid-coronal slice – volumetric ROI<br><br>Basal ganglia – volumetric ROI |
| **Corpus callosum;** | Corpus Callosum: genu, body and splenium, - sagittal plane ROI at mid-sagittal slice<br><br>White matter on outer edge of the medial frontal, parietal and occipital lobes – volumetric ROI | Basal ganglia, midbrain and brainstem – volumetric ROI<br><br>White matter on outer edge of the lateral frontal, parietal and occipital lobes – volumetric ROI |
| **Genu** | Genu of the corpus callosum - sagittal plane ROI at mid-sagittal slice | Medial and posterior frontal lobe – coronal plane ROI |

| | | |
|---|---|---|
| **Body** | Body of the corpus callosum - sagittal plane ROI at mid-sagittal slice | Anterior frontal lobe – coronal plane ROI<br><br>Posterior parietal and occipital lobes – coronal plane ROI |
| **Splenium** | Splenium of the corpus callosum - sagittal plane ROI at mid-sagittal slice | Anterior parietal lobe – coronal plane ROI |

**Supplementary Table 2: Inclusion and exclusion regions of interest used in tractography.** Anatomical locations of regions of interests (ROIs) utilized to extract major white matter tracts from the whole-brain tractogram. ROIs were either 2D slices drawn in sagittal, axial or coronal view, or 3D volumetric ROIs. For segments of the corpus callosum, the corpus callosum was first extracted in full, then subsequently segmented into the genu, body and splenium using more spatially specific ROIs.

| | Rate of change (Δ/year) | Predicted value at age 4 | Predicted value at age 8 | Percent increase from ages 4-8 |
|---|---|---|---|---|
| Left Arcuate Fasciculus | 5.064E-03* | 0.300 | 0.320 | 6.748 |
| Right Arcuate Fasciculus | 4.689E-03* | 0.309 | 0.328 | 6.062 |
| Corpus Callosum Body | 3.531E-03* | 0.457 | 0.471 | 3.089 |
| Corpus Callosum Genu | -2.528E-04 | 0.377 | 0.376 | -0.268 |
| Corpus Callosum Splenium | 7.060E-03* | 0.399 | 0.427 | 7.078 |
| Left Cingulum | 4.185E-03* | 0.328 | 0.344 | 5.110 |
| Right Cingulum | 5.053E-03* | 0.324 | 0.344 | 6.238 |
| Left Corticospinal Tract | 7.719E-03* | 0.513 | 0.544 | 6.019 |
| Right Corticospinal Tract | 7.165E-03* | 0.499 | 0.527 | 5.747 |
| Left Fornix | 6.184E-03 | 0.461 | 0.486 | 5.360 |
| Right Fornix | 4.431E-03 | 0.455 | 0.472 | 3.899 |
| Left Inf. Frontal-Occipital Fasciculus | 2.445E-03 | 0.445 | 0.455 | 2.197 |
| Right Inf. Frontal-Occipital Fasciculus | 4.753E-03* | 0.448 | 0.467 | 4.241 |
| Left Inf. Longitudinal Fasciculus | 3.094E-03 | 0.335 | 0.347 | 3.695 |
| Right Inf. Longitudinal Fasciculus | 1.843E-03 | 0.343 | 0.351 | 2.148 |
| Left Sup. Longitudinal Fasciculus | -5.815E-04 | 0.295 | 0.292 | -0.790 |
| Right Sup. Longitudinal Fasciculus | -1.661E-04 | 0.298 | 0.298 | -0.223 |
| Left Uncinate Fasciculus | -6.502E-04 | 0.377 | 0.375 | -0.689 |
| Right Uncinate Fasciculus | -5.736E-04 | 0.373 | 0.371 | -0.615 |
| Whole Brain | 2.552E-03* | 0.309 | 0.319 | 3.304 |

**Supplementary Table 3: Developmental trajectory values for fiber density.** * indicates tracts for which there was a significant longitudinal increase from ages 4-8.

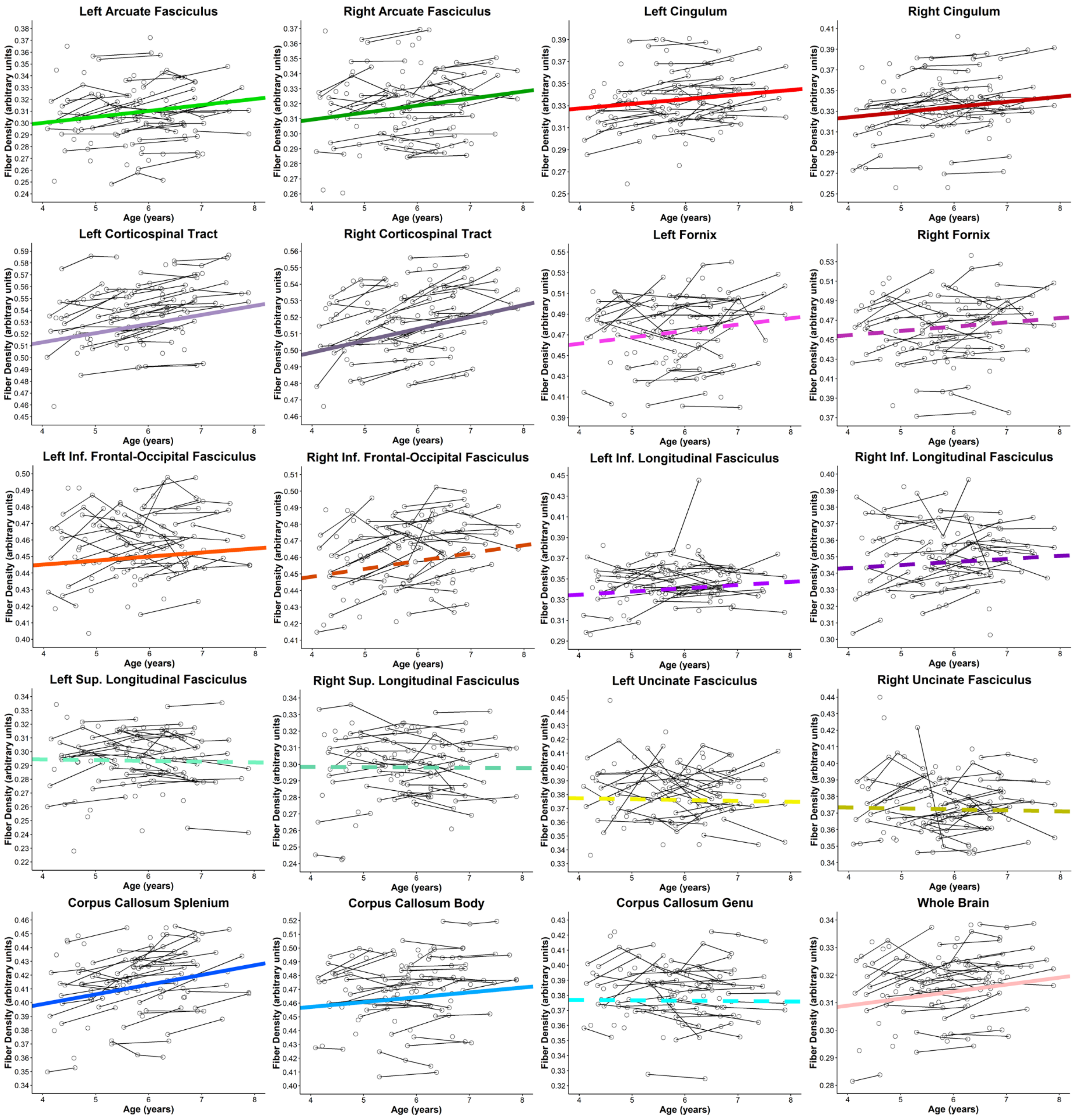


**Supplementary Figure 1: Scatterplots of tract-wise fiber density from 4 to 8 years of age.** Scatterplots of tract-wise FD values for individual scans. Open circles indicate individual scan values. Scans collected from the same participant are indicated by a black line connecting the open circles. The bold coloured lines indicate the average linear trend within each imaging metric. Solid lines indicate tracts that had significant change from ages 4 to 8, while dashed lines indicate tracts with no significant change.

| | Rate of change (Δ/year) | Predicted value at age 4 | Predicted value at age 8 | Percent increase from ages 4-8 |
|---|---|---|---|---|
| Left Arcuate Fasciculus | 1.780E-02* | 1.058 | 1.129 | 6.729 |
| Right Arcuate Fasciculus | 2.165E-02* | 1.047 | 1.134 | 8.271 |
| Corpus Callosum Body | 2.032E-02* | 1.026 | 1.107 | 7.926 |
| Corpus Callosum Genu | -1.823E-03 | 1.089 | 1.082 | -0.670 |
| Corpus Callosum Splenium | 2.205E-02* | 1.043 | 1.131 | 8.457 |
| Left Cingulum | 1.144E-02* | 0.981 | 1.027 | 4.662 |
| Right Cingulum | 1.062E-02* | 0.985 | 1.028 | 4.310 |
| Left Corticospinal Tract | 3.176E-02* | 0.990 | 1.117 | 12.830 |
| Right Corticospinal Tract | 3.213E-02* | 0.991 | 1.120 | 12.967 |
| Left Fornix | 1.539E-02* | 0.992 | 1.053 | 6.207 |
| Right Fornix | 1.404E-02* | 0.992 | 1.048 | 5.662 |
| Left Inf. Frontal-Occipital Fasciculus | 1.637E-02* | 1.068 | 1.133 | 6.132 |
| Right Inf. Frontal-Occipital Fasciculus | 2.105E-02* | 1.056 | 1.140 | 7.970 |
| Left Inf. Longitudinal Fasciculus | 3.451E-02* | 1.140 | 1.278 | 12.104 |
| Right Inf. Longitudinal Fasciculus | 3.492E-02* | 1.155 | 1.294 | 12.099 |
| Left Sup. Longitudinal Fasciculus | 1.779E-02* | 1.088 | 1.159 | 6.542 |
| Right Sup. Longitudinal Fasciculus | 1.865E-02* | 1.083 | 1.157 | 6.888 |
| Left Uncinate Fasciculus | 1.466E-02* | 1.124 | 1.183 | 5.218 |
| Right Uncinate Fasciculus | 1.085E-02* | 1.121 | 1.165 | 3.871 |
| Whole Brain | 1.515E-02* | 1.055 | 1.115 | 5.745 |

**Supplementary Table 4: Developmental trajectory values for fiber cross-section.** * indicates tracts for which there was a significant change from ages 4-8.

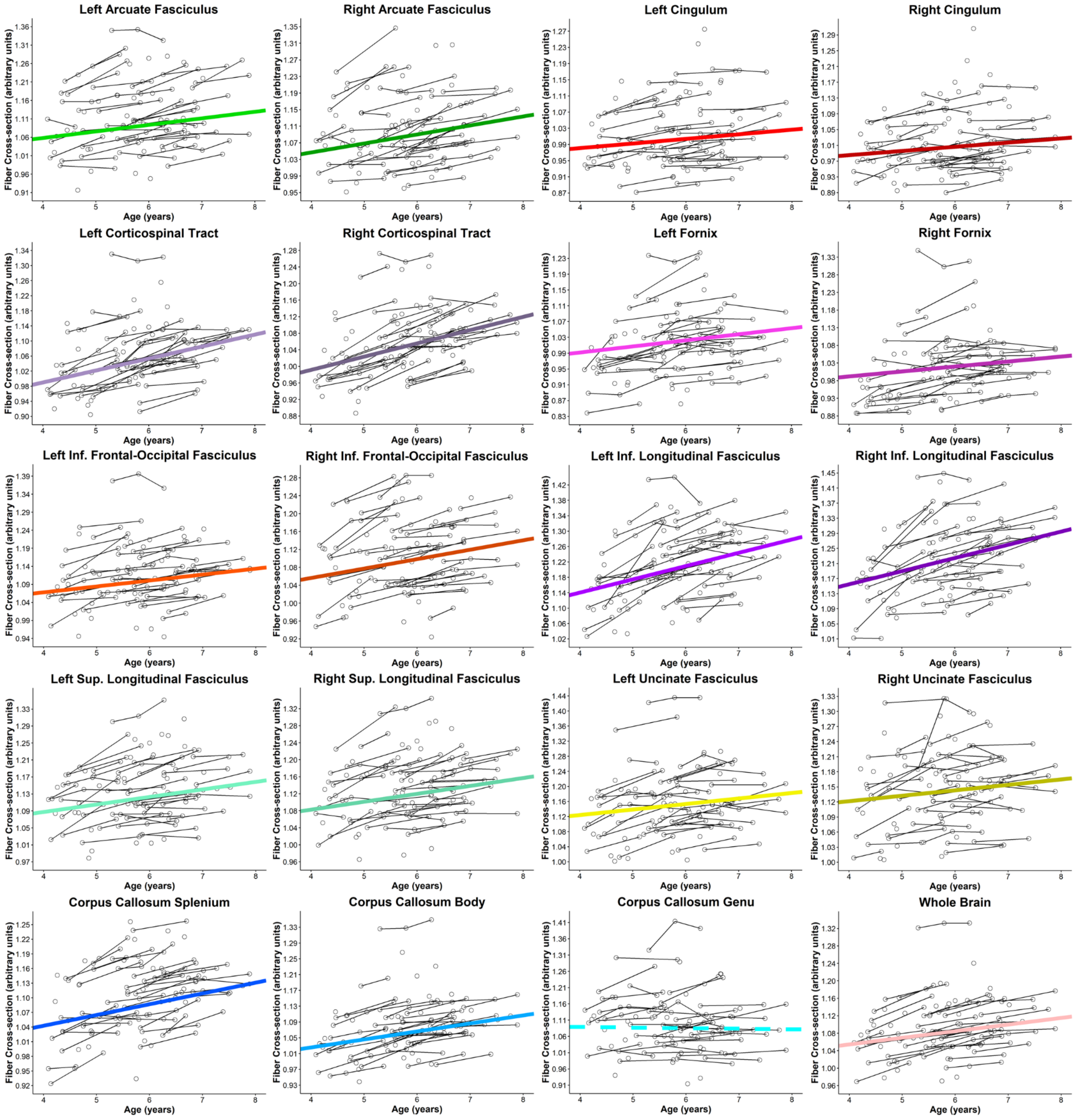


**Supplementary Figure 2: Scatterplots of tract-wise fiber cross-section from 4 to 8 years of age.** Scatterplots of tract-wise FC values for individual scans. Open circles indicate individual scan values. Scans collected from the same participant are indicated by a black line connecting the open circles. The bold coloured lines indicate the average linear trend within each imaging metric. Solid lines indicate tracts that had significant change from ages 4 to 8, while dashed lines indicate tracts with no significant change.

| | Rate of change ($\Delta$/year) | Predicted value at age 4 | Predicted value at age 8 | Percent increase from ages 4-8 |
|---|---|---|---|---|
| Left Arcuate Fasciculus | 1.129E-02* | 0.319 | 0.364 | 14.147 |
| Right Arcuate Fasciculus | 1.297E-02* | 0.324 | 0.376 | 16.023 |
| Corpus Callosum Body | 1.411E-02* | 0.463 | 0.519 | 12.189 |
| Corpus Callosum Genu | -4.172E-04 | 0.405 | 0.404 | -0.412 |
| Corpus Callosum Splenium | 1.827E-02* | 0.406 | 0.479 | 17.985 |
| Left Cingulum | 8.341E-03* | 0.322 | 0.355 | 10.372 |
| Right Cingulum | 9.514E-03* | 0.318 | 0.356 | 11.956 |
| Left Corticospinal Tract | 2.682E-02* | 0.503 | 0.610 | 21.325 |
| Right Corticospinal Tract | 2.626E-02* | 0.487 | 0.592 | 21.563 |
| Left Fornix | 1.332E-02* | 0.455 | 0.509 | 11.702 |
| Right Fornix | 1.128E-02* | 0.447 | 0.492 | 10.086 |
| Left Inf. Frontal-Occipital Fasciculus | 1.016E-02* | 0.472 | 0.512 | 8.615 |
| Right Inf. Frontal-Occipital Fasciculus | 1.459E-02* | 0.470 | 0.528 | 12.425 |
| Left Inf. Longitudinal Fasciculus | 1.670E-02* | 0.383 | 0.450 | 17.434 |
| Right Inf. Longitudinal Fasciculus | 1.478E-02* | 0.398 | 0.457 | 14.854 |
| Left Sup. Longitudinal Fasciculus | 4.827E-03 | 0.321 | 0.341 | 6.008 |
| Right Sup. Longitudinal Fasciculus | 5.641E-03* | 0.323 | 0.346 | 6.983 |
| Left Uncinate Fasciculus | 4.388E-03 | 0.423 | 0.441 | 4.148 |
| Right Uncinate Fasciculus | 3.750E-03 | 0.416 | 0.431 | 3.608 |
| Whole Brain | 8.655E-03* | 0.323 | 0.358 | 10.713 |

**Supplementary Table 5: Developmental trajectory values for fiber density and cross-section.** * indicates tracts for which there was a significant change from ages 4-8.

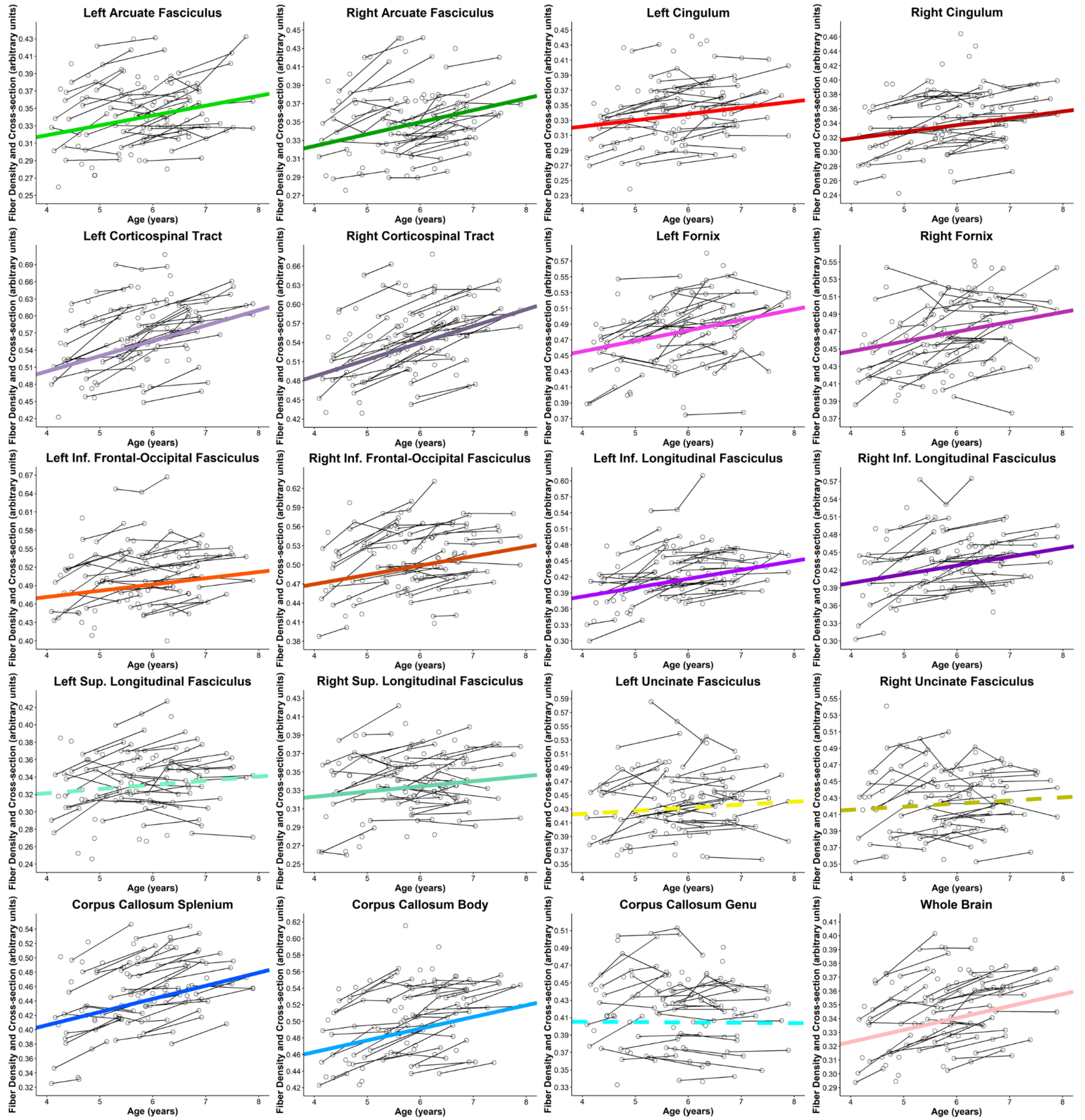


**Supplementary Figure 3: Scatterplots of tract-wise fiber density and cross-section from 4 to 8 years of age.** Scatterplots of tract-wise FDC values for individual scans. Open circles indicate individual scan values. Scans collected from the same participant are indicated by a black line connecting the open circles. The bold coloured lines indicate the average linear trend within each imaging metric. Solid lines indicate tracts that had significant change from ages 4 to 8, while dashed lines indicate tracts with no significant change.

| | Rate of change ($\Delta$/year) | Predicted value at age 4 | Predicted value at age 8 | Percent increase from ages 4-8 |
|---|---|---|---|---|
| Left Arcuate Fasciculus | 5.45E-03* | 0.256 | 0.278 | 7.852 |
| Right Arcuate Fasciculus | 5.26E-03* | 0.265 | 0.286 | 7.358 |
| Corpus Callosum Body | 4.77E-03* | 0.291 | 0.310 | 6.152 |
| Corpus Callosum Genu | 6.26E-04 | 0.270 | 0.272 | 0.919 |
| Corpus Callosum Splenium | 4.49E-03* | 0.276 | 0.294 | 6.106 |
| Left Cingulum | 1.30E-03 | 0.260 | 0.265 | 1.957 |
| Right Cingulum | 3.81E-03 | 0.245 | 0.260 | 5.859 |
| Left Corticospinal Tract | 6.79E-03* | 0.326 | 0.353 | 7.690 |
| Right Corticospinal Tract | 5.24E-03* | 0.331 | 0.352 | 5.952 |
| Left Fornix | 3.21E-03 | 0.257 | 0.270 | 4.754 |
| Right Fornix | 3.05E-03 | 0.259 | 0.271 | 4.502 |
| Left Inf. Frontal-Occipital Fasciculus | 4.21E-03* | 0.301 | 0.317 | 5.307 |
| Right Inf. Frontal-Occipital Fasciculus | 4.13E-03* | 0.309 | 0.325 | 5.073 |
| Left Inf. Longitudinal Fasciculus | 2.44E-03 | 0.221 | 0.231 | 4.226 |
| Right Inf. Longitudinal Fasciculus | 2.03E-03 | 0.235 | 0.243 | 3.347 |
| Left Sup. Longitudinal Fasciculus | 5.60E-03* | 0.310 | 0.332 | 6.744 |
| Right Sup. Longitudinal Fasciculus | 5.16E-03* | 0.325 | 0.346 | 5.976 |
| Left Uncinate Fasciculus | 2.20E-03 | 0.236 | 0.245 | 3.587 |
| Right Uncinate Fasciculus | 2.63E-04 | 0.244 | 0.245 | 0.429 |
| Whole Brain | 2.37E-03* | 0.222 | 0.231 | 4.094 |

**Supplementary Table 6: Developmental trajectory values for fractional anisotropy.** * indicates tracts for which there was a significant change from ages 4-8.

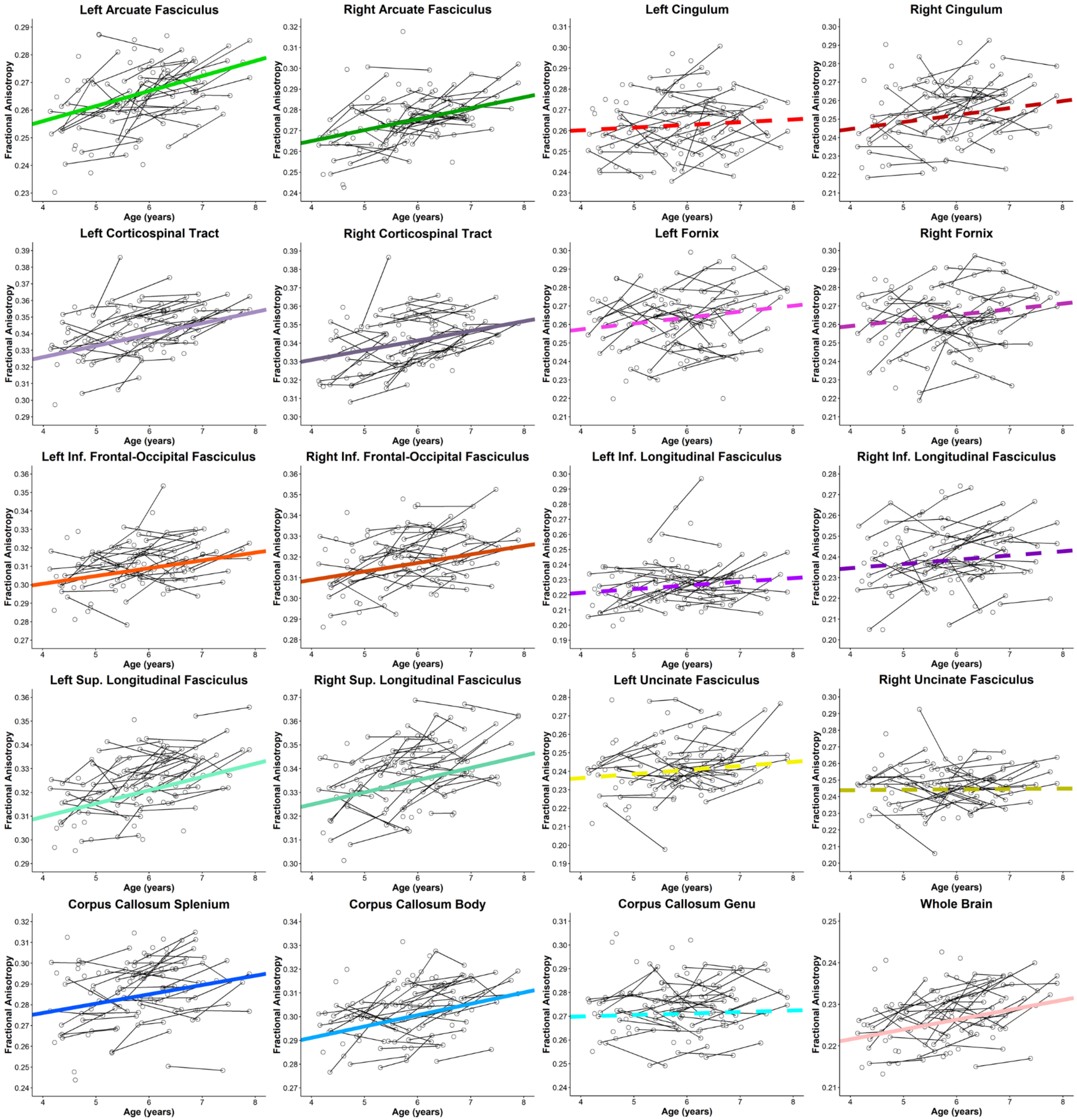


**Supplementary Figure 4: Scatterplots of tract-wise fractional anisotropy from 4 to 8 years of age.** Scatterplots of tract-wise FA values for individual scans. Open circles indicate individual scan values. Scans collected from the same participant are indicated by a black line connecting the open circles. The bold coloured lines indicate the average linear trend within each imaging metric. Solid lines indicate tracts that had significant change from ages 4 to 8, while dashed lines indicate tracts with no significant change.

| | Rate of change (Δ/year) | Predicted value at age 4 | Predicted value at age 8 | Percent decrease from ages 4-8 |
|---|---|---|---|---|
| Left Arcuate Fasciculus | -1.045E-05* | 8.458E-04 | 8.040E-04 | 5.200 |
| Right Arcuate Fasciculus | -8.998E-06* | 8.067E-04 | 7.707E-04 | 4.670 |
| Corpus Callosum Body | -1.144E-05* | 9.169E-04 | 8.712E-04 | 5.252 |
| Corpus Callosum Genu | -9.068E-06* | 9.546E-04 | 9.183E-04 | 3.950 |
| Corpus Callosum Splenium | -1.258E-05* | 9.066E-04 | 8.563E-04 | 5.878 |
| Left Cingulum | -8.510E-06* | 8.627E-04 | 8.287E-04 | 4.108 |
| Right Cingulum | -9.601E-06* | 8.582E-04 | 8.198E-04 | 4.685 |
| Left Corticospinal Tract | -1.145E-05* | 8.504E-04 | 8.046E-04 | 5.693 |
| Right Corticospinal Tract | -9.641E-06* | 8.356E-04 | 7.970E-04 | 4.839 |
| Left Fornix | -9.901E-06 | 1.247E-03 | 1.207E-03 | 3.281 |
| Right Fornix | -1.148E-05 | 1.235E-03 | 1.189E-03 | 3.864 |
| Left Inf. Frontal-Occipital Fasciculus | -1.053E-05* | 8.715E-04 | 8.294E-04 | 5.079 |
| Right Inf. Frontal-Occipital Fasciculus | -1.067E-05* | 8.563E-04 | 8.137E-04 | 5.244 |
| Left Inf. Longitudinal Fasciculus | -9.684E-06* | 8.665E-04 | 8.278E-04 | 4.680 |
| Right Inf. Longitudinal Fasciculus | -1.087E-05* | 8.419E-04 | 7.984E-04 | 5.444 |
| Left Sup. Longitudinal Fasciculus | -1.169E-05* | 8.389E-04 | 7.921E-04 | 5.905 |
| Right Sup. Longitudinal Fasciculus | -1.146E-05* | 8.182E-04 | 7.724E-04 | 5.935 |
| Left Uncinate Fasciculus | -1.013E-05* | 9.060E-04 | 8.655E-04 | 4.683 |
| Right Uncinate Fasciculus | -9.066E-06* | 8.798E-04 | 8.435E-04 | 4.299 |
| Whole Brain | -8.634E-06* | 9.180E-04 | 8.835E-04 | 3.909 |

**Supplementary Table 7: Developmental trajectory values for mean diffusivity.** * indicates tracts for which there was a significant change from ages 4-8.

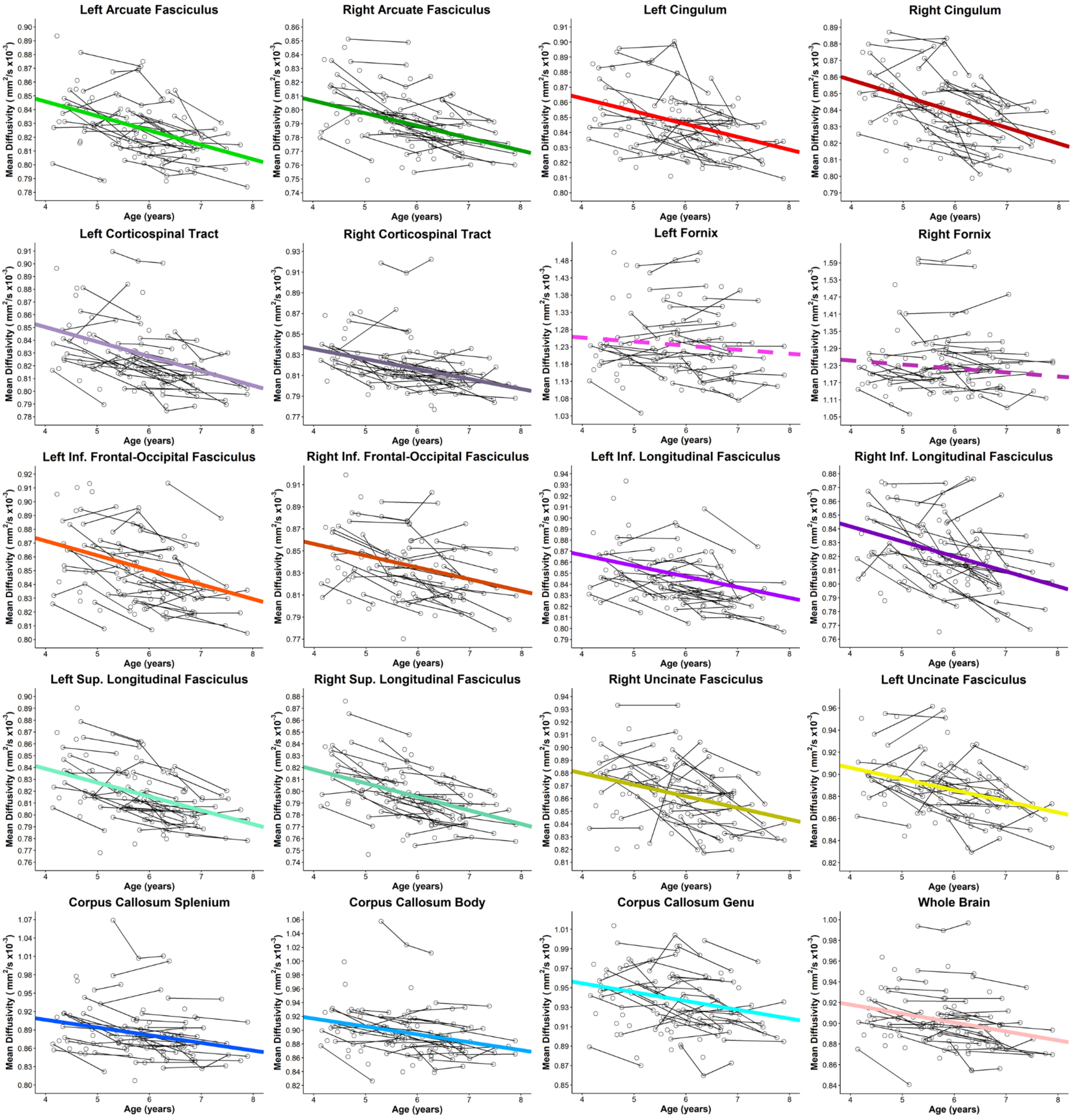


**Supplementary Figure 5: Scatterplots of tract-wise mean diffusivity from 4 to 8 years of age.** Scatterplots of tract-wise MD values for individual scans. Open circles indicate individual scan values. Scans collected from the same participant are indicated by a black line connecting the open circles. The bold coloured lines indicate the average linear trend within each imaging metric. Solid lines indicate tracts that had significant change from ages 4 to 8, while dashed lines indicate tracts with no significant change.

| | Rate of change ($\Delta$/year) | Predicted value at age 4 | Predicted value at age 8 | Percent decrease from ages 4-8 |
|---|---|---|---|---|
| Left Arcuate Fasciculus | -1.142E-05* | 7.333E-04 | 6.876E-04 | 6.643 |
| Right Arcuate Fasciculus | -1.002E-05* | 6.965E-04 | 6.564E-04 | 6.106 |
| Corpus Callosum Body | -1.196E-05* | 7.646E-04 | 7.167E-04 | 6.674 |
| Corpus Callosum Genu | -7.911E-06* | 8.090E-04 | 7.774E-04 | 4.070 |
| Corpus Callosum Splenium | -1.299E-05* | 7.597E-04 | 7.077E-04 | 7.344 |
| Left Cingulum | -7.653E-06* | 7.406E-04 | 7.100E-04 | 4.311 |
| Right Cingulum | -1.000E-05* | 7.461E-04 | 7.061E-04 | 5.665 |
| Left Corticospinal Tract | -1.292E-05* | 6.959E-04 | 6.442E-04 | 8.022 |
| Right Corticospinal Tract | -1.062E-05* | 6.812E-04 | 6.388E-04 | 6.651 |
| Left Fornix | -1.048E-05 | 1.071E-03 | 1.029E-03 | 4.072 |
| Right Fornix | -1.206E-05 | 1.064E-03 | 1.016E-03 | 4.748 |
| Left Inf. Frontal-Occipital Fasciculus | -1.099E-05* | 7.270E-04 | 6.830E-04 | 6.437 |
| Right Inf. Frontal-Occipital Fasciculus | -1.096E-05* | 7.102E-04 | 6.664E-04 | 6.578 |
| Left Inf. Longitudinal Fasciculus | -9.411E-06* | 7.619E-04 | 7.242E-04 | 5.198 |
| Right Inf. Longitudinal Fasciculus | -1.041E-05* | 7.347E-04 | 6.930E-04 | 6.009 |
| Left Sup. Longitudinal Fasciculus | -1.233E-05* | 6.982E-04 | 6.489E-04 | 7.600 |
| Right Sup. Longitudinal Fasciculus | -1.189E-05* | 6.719E-04 | 6.243E-04 | 7.621 |
| Left Uncinate Fasciculus | -9.705E-06* | 7.904E-04 | 7.516E-04 | 5.165 |
| Right Uncinate Fasciculus | -7.989E-06* | 7.638E-04 | 7.318E-04 | 4.367 |
| Whole Brain | -8.550E-06* | 8.082E-04 | 7.740E-04 | 4.418 |

**Supplementary Table 8: Developmental trajectory values for radial diffusivity.** * indicates tracts for which there was a significant change from ages 4-8.

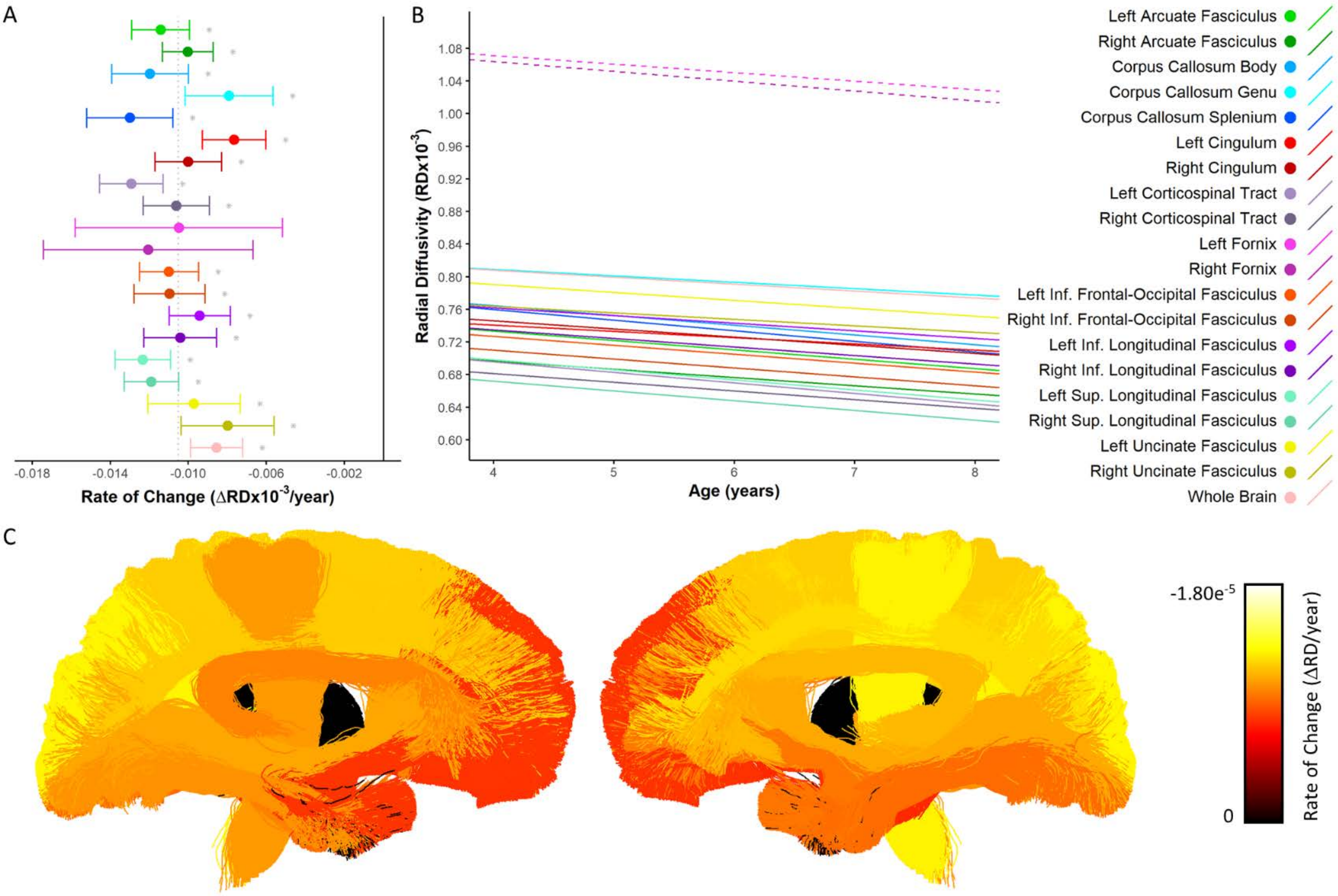


**Supplementary Figure 6. Developmental changes in radial diffusivity.** A) Rate of change in RD shown for each tract. * denotes tracts in which there was significant growth. The solid black vertical line indicates zero change, while the dotted gray vertical line indicates the mean rate of change across all tracts. B) Linear trajectories of RD growth for each tract. Solid lines indicate tracts in which there was a statistically significant relationship with age; dotted lines correspond to non-significant tracts. C) Anatomical visualization of tract-wise rates of change in RD. Streamlines corresponding to each tract are coloured according to the rate of change of RD observed in that tract.

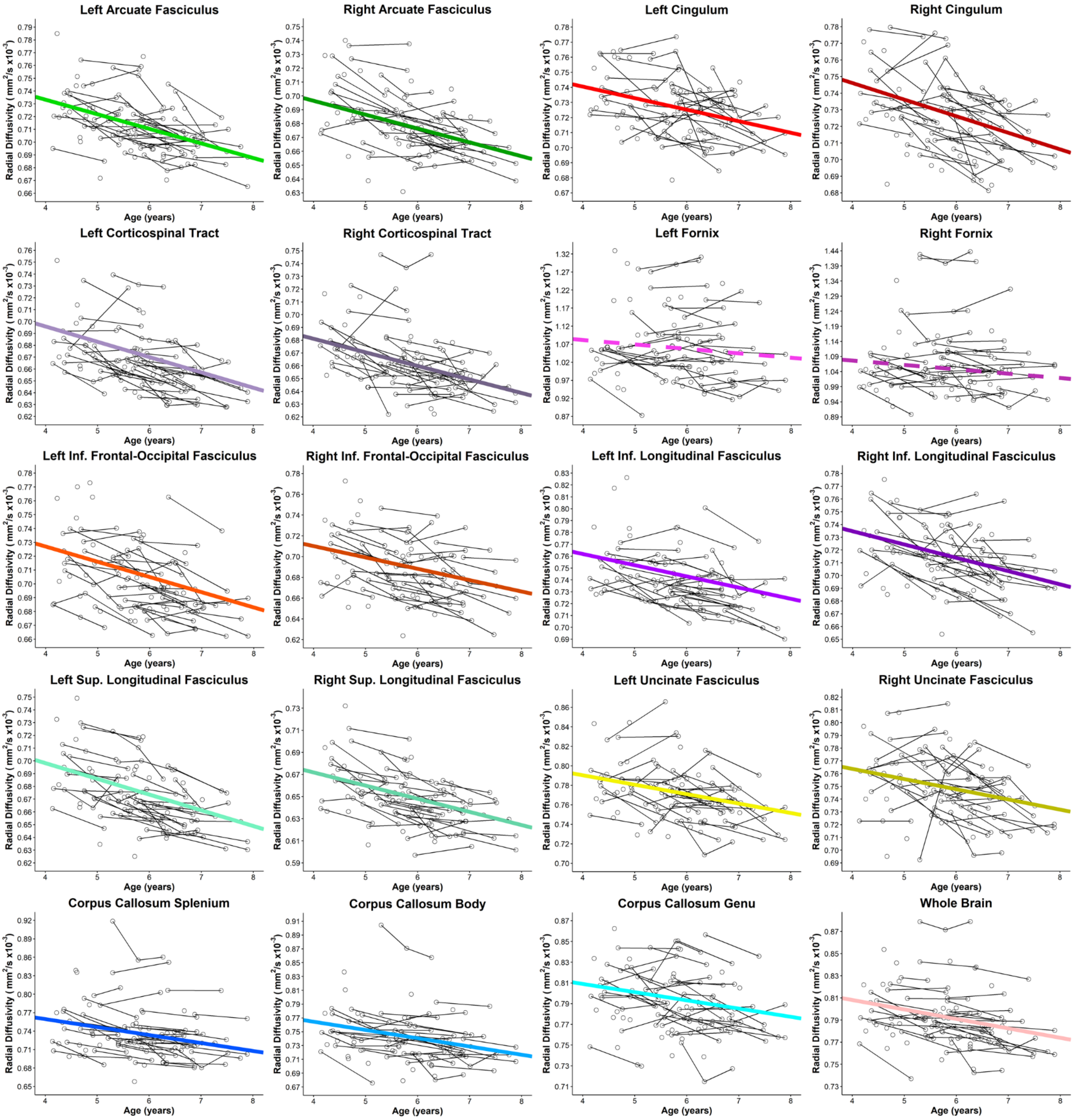


**Supplementary Figure 7: Scatterplots of tract-wise radial diffusivity from 4 to 8 years of age.** Scatterplots of tract-wise RD values for individual scans. Open circles indicate individual scan values. Scans collected from the same participant are indicated by a black line connecting the open circles. The bold coloured lines indicate the average linear trend within each imaging metric. Solid lines indicate tracts that had significant change from ages 4 to 8, while dashed lines indicate tracts with no significant change.

| | Rate of change ($\Delta$/year) | Predicted value at age 4 | Predicted value at age 8 | Percent decrease from ages 4-8 |
|---|---|---|---|---|
| Left Arcuate Fasciculus | -8.642E-06* | 1.071E-03 | 1.037E-03 | 3.33475 |
| Right Arcuate Fasciculus | -7.163E-06* | 1.027E-03 | 9.988E-04 | 2.868532 |
| Corpus Callosum Body | -1.050E-05* | 1.222E-03 | 1.180E-03 | 3.560873 |
| Corpus Callosum Genu | -1.149E-05* | 1.246E-03 | 1.200E-03 | 3.829754 |
| Corpus Callosum Splenium | -1.175E-05* | 1.200E-03 | 1.153E-03 | 4.075244 |
| Left Cingulum | -1.025E-05* | 1.107E-03 | 1.066E-03 | 3.845947 |
| Right Cingulum | -8.760E-06* | 1.082E-03 | 1.047E-03 | 3.34617 |
| Left Corticospinal Tract | -8.715E-06* | 1.160E-03 | 1.125E-03 | 3.09883 |
| Right Corticospinal Tract | -7.570E-06* | 1.144E-03 | 1.114E-03 | 2.71919 |
| Left Fornix | -1.055E-05 | 1.608E-03 | 1.566E-03 | 2.695647 |
| Right Fornix | -1.023E-05 | 1.576E-03 | 1.535E-03 | 2.665572 |
| Left Inf. Frontal-Occipital Fasciculus | -9.943E-06* | 1.161E-03 | 1.121E-03 | 3.546377 |
| Right Inf. Frontal-Occipital Fasciculus | -9.975E-06* | 1.149E-03 | 1.109E-03 | 3.598945 |
| Left Inf. Longitudinal Fasciculus | -1.072E-05* | 1.077E-03 | 1.034E-03 | 4.149169 |
| Right Inf. Longitudinal Fasciculus | -1.189E-05* | 1.057E-03 | 1.009E-03 | 4.713464 |
| Left Sup. Longitudinal Fasciculus | -1.037E-05* | 1.120E-03 | 1.079E-03 | 3.843648 |
| Right Sup. Longitudinal Fasciculus | -1.091E-05* | 1.111E-03 | 1.068E-03 | 4.085915 |
| Left Uncinate Fasciculus | -1.113E-05* | 1.137E-03 | 1.093E-03 | 4.072709 |
| Right Uncinate Fasciculus | -1.127E-05* | 1.112E-03 | 1.067E-03 | 4.226024 |
| Whole Brain | -9.029E-06* | 1.138E-03 | 1.102E-03 | 3.277313 |

**Supplementary Table 9: Developmental trajectory values for axial diffusivity.** * indicates tracts for which there was a significant change from ages 4-8.

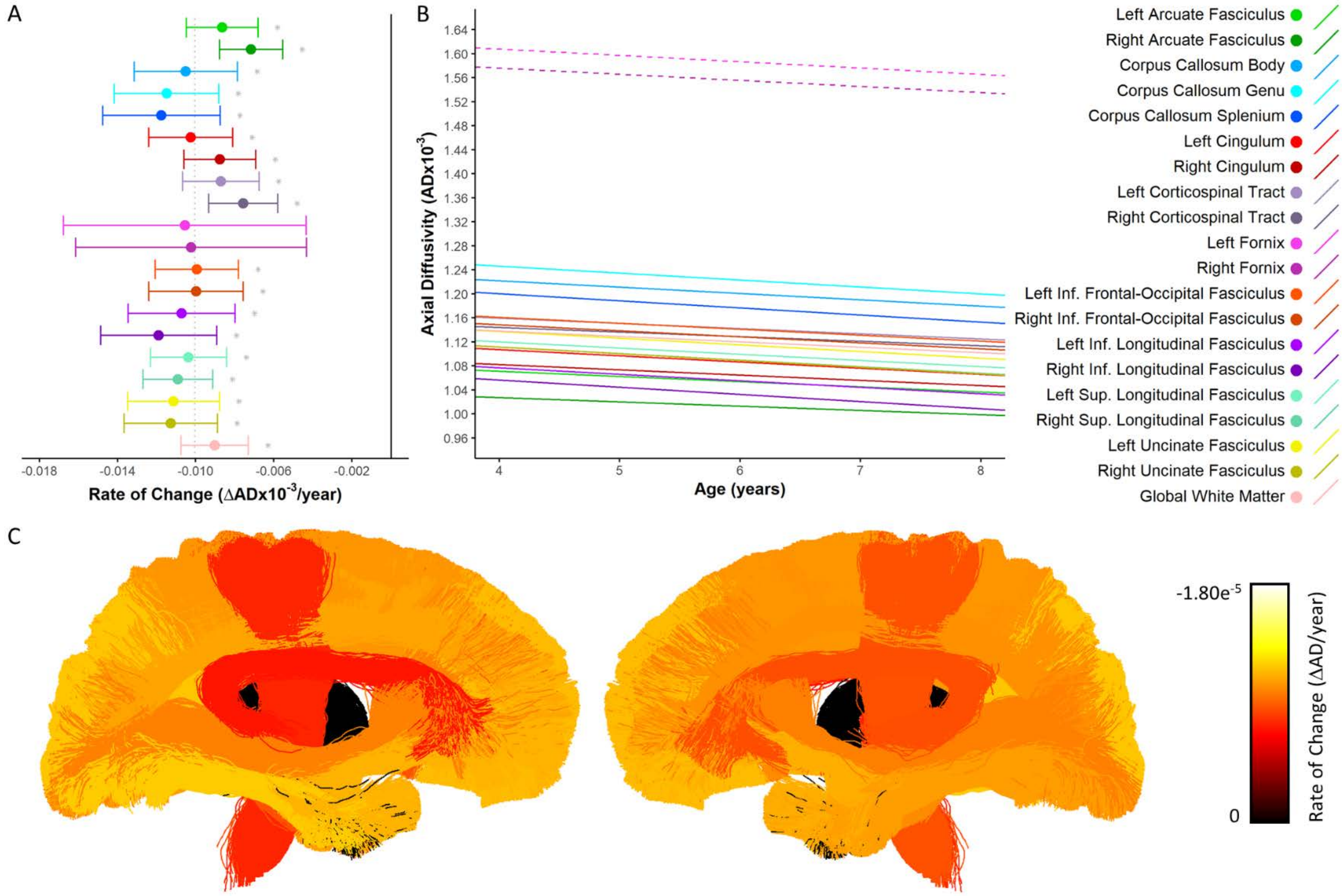


**Supplementary Figure 8. Developmental changes in axial diffusivity.** A) Rate of change in AD shown for each tract. * denotes tracts in which there was significant growth. The solid black vertical line indicates zero change, while the dotted gray vertical line indicates the mean rate of change across all tracts. B) Linear trajectories of AD growth for each tract. Solid lines indicate tracts in which there was a statistically significant relationship with age; dotted lines correspond to non-significant tracts. C) Anatomical visualization of tract-wise rates of change in AD. Streamlines corresponding to each tract are coloured according to the rate of change of AD observed in that tract.

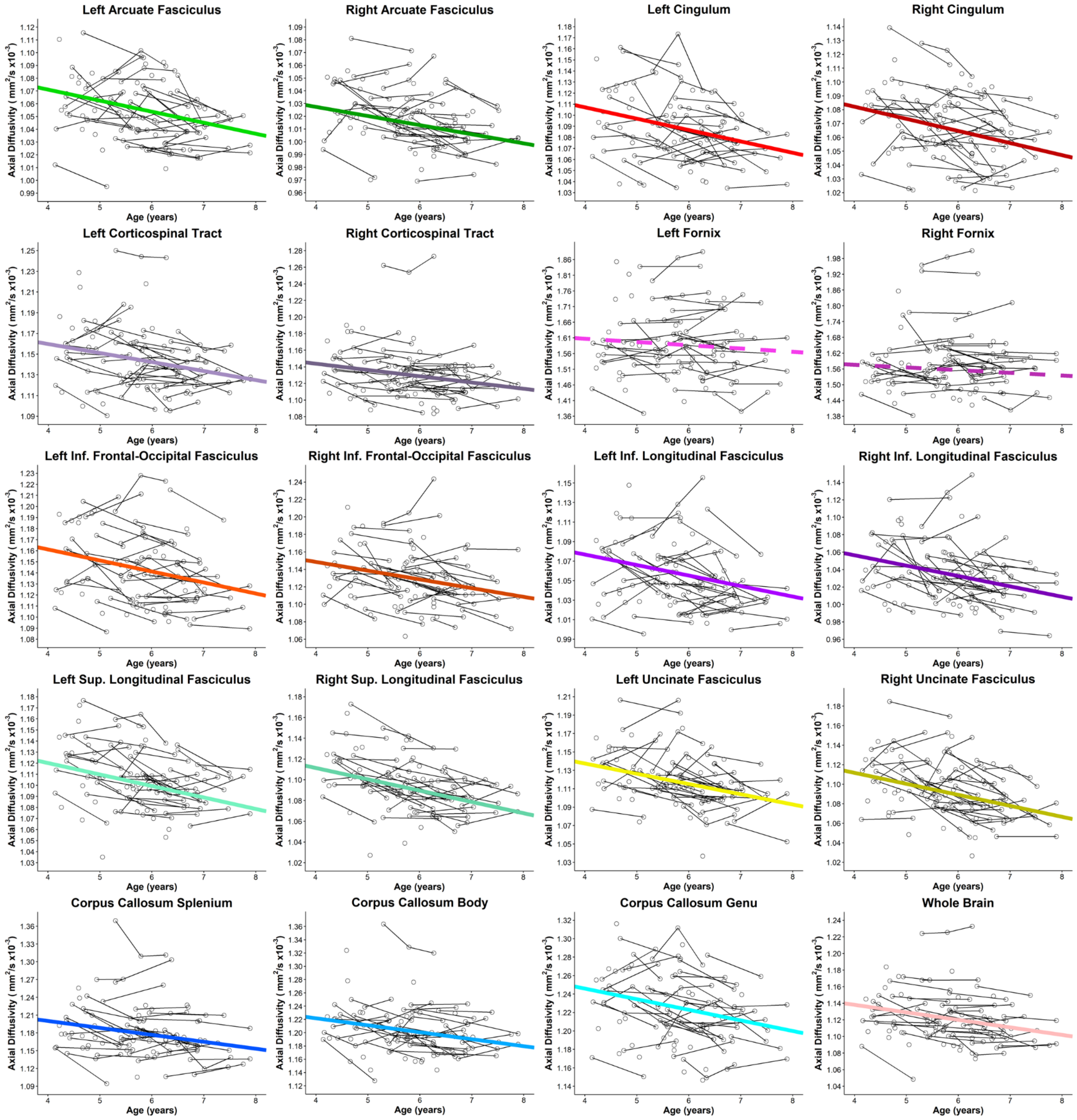


**Supplementary Figure 9: Scatterplots of tract-wise axial diffusivity from 4 to 8 years of age.** Scatterplots of tract-wise AD values for individual scans. Open circles indicate individual scan values. Scans collected from the same participant are indicated by a black line connecting the open circles. The bold coloured lines indicate the average linear trend within each imaging metric. Solid lines indicate tracts that had significant change from ages 4 to 8, while dashed lines indicate tracts with no significant change.

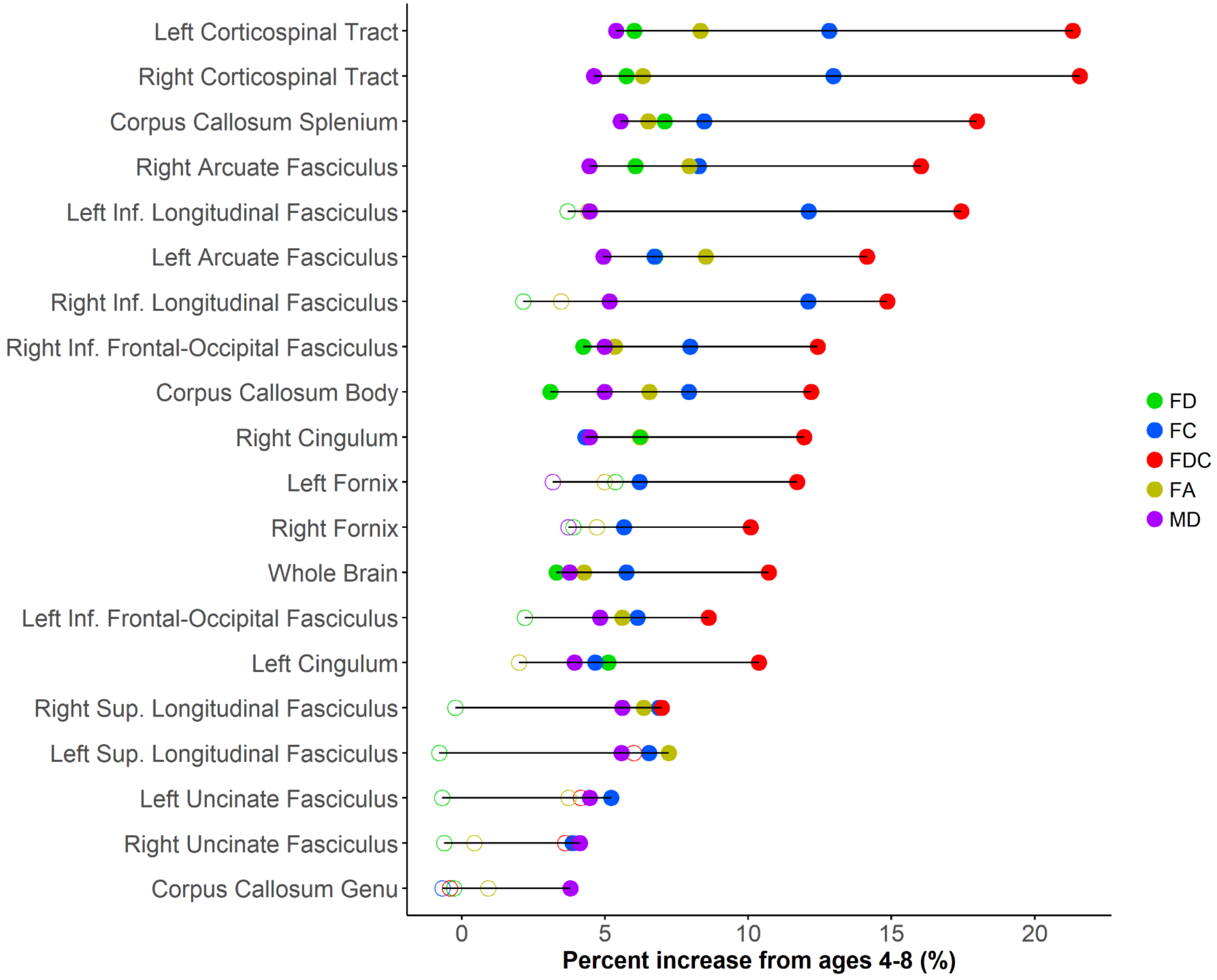


**Supplementary Figure 10: Percent increase in imaging metrics from 4 to 8 years of age.** Percent increase in FD, FC, FDC, FA and MD from ages 4-8. Tracts are listed in descending order of average percent change calculated across all included metrics. Solid circles indicate statistically significant growth, while open circles indicate non-significant growth.

| | **FD** | **FC** | **FDC** | **FA** | **MD** | **RD** | **AD** |
|---|---|---|---|---|---|---|---|
| **FD** | 1 | 0.102 | .635* | .576* | -.746* | -.780* | -.610* |
| **FC** | 0.102 | 1 | .830* | .523* | 0.163 | 0.096 | 0.100 |
| **FDC** | .635* | .830* | 1 | .724* | -.291* | -.360* | -0.298* |
| **FA** | .576* | .523* | .724* | 1 | -.413* | -.507* | -.306* |
| **MD** | -.746* | 0.163 | -.291* | -.413* | 1 | .994* | .980* |
| **RD** | -.780* | 0.096 | -.360* | -.507* | .994* | 1 | .952* |
| **AD** | -.665* | 0.100 | -0.298* | -.306* | .980* | .952* | 1 |

**Supplementary Table 10: Correlations between imaging metrics of the global white matter mask.** * indicates significant correlations at FDR-corrected p-value<0.05.

| | **ΔFD** | **ΔFC** | **ΔFDC** | **ΔFA** | **ΔMD** | **ΔRD** | **ΔAD** |
|---|---|---|---|---|---|---|---|
| **ΔFD** | 1 | .585* | .900* | -0.017 | -.695* | -.642* | -.629* |
| **ΔFC** | .585* | 1 | .868* | -0.119 | -0.239 | -0.185 | -0.268 |
| **ΔFDC** | .900* | .868* | 1 | -0.087 | -.522* | -.458* | -.509* |
| **ΔFA** | -0.017 | -0.119 | -0.087 | 1 | -0.347* | -.387* | .407* |
| **ΔMD** | -.695* | -0.239 | -.522* | -0.074 | 1 | .947* | .877* |
| **ΔRD** | -.642* | -0.185 | -.458* | -.387* | .947* | 1 | .676* |
| **ΔAD** | -.629* | -0.268 | -.509* | .407* | .877* | .676* | 1 |

**Supplementary Table 11: Correlations between longitudinal changes in imaging metrics of the global white matter mask.** * indicates significant correlations at FDR-corrected p-value<0.05.
